# *Osiris* gene family defines the cuticle nano-patterns of *Drosophila*

**DOI:** 10.1101/2024.02.06.579014

**Authors:** Zhengkuan Sun, Sachi Inagaki, Keita Miyoshi, Kuniaki Saito, Shigeo Hayashi

## Abstract

Nanostructures of pores and protrusions in the insect cuticle modify molecular permeability and surface wetting and help insects sense a variety of environmental cues. The cellular mechanism specifying cuticle nanostructures is poorly understood. Here, we show that insect-specific *Osiris* family genes are expressed in various cuticle-secreting cells in the *Drosophila* head in the early stage of cuticle secretion and collectively cover nearly the entire surface of the head epidermis. We show that each sense organ cell with various cuticular nanostructures expresses a unique combination of *Osiris* genes. *Osiris* gene mutations caused various cuticle defects in the corneal nipples of the eye and pores of the chemosensory sensilla. *Osiris* genes provide an entry point for investigating cuticle nanopatterning in insects.

## 1. Introduction

Extracellular materials cover the body surface of every higher animal and plant in the form of stratum corneum, cell wall, or cuticle, protecting fragile internal body environments from the external world filled with toxic chemicals, genotoxic radiations, and predators. Those extracellular matrices are denucleated remnants of keratinocytes of the vertebrate epidermis or the cellulose-based plant cell walls.

In insects, cuticles are multilayered structures consisting of chitin-rich procuticles covered by protein and lipid-rich epicuticles, secreted sequentially by the epidermal cells in an outside-to-inside-order (Wigglesworth 1948). The cuticles harden to form protective shells that serve as exoskeletons. In addition, cuticles of sensory organs serve as the window to receive environmental signals such as light, chemicals, and mechanical stimuli (Stocker 1994). The insect sensillum comprises the hair (bristle) and socket cuticles, each secreted from trichogen and tormogen cells (Shanbhag *et al*. 1999). Sensory neurons innervate inside the hair cell cuticles and are associated with glia and sheath cells. All cells in each sensillum are descendants of single sensory precursor cells uniquely fated for specific sensory lineage (Hartenstein and Posakony 1989).

Cuticles of sense organs adopt specific nanostructures to optimize the reception of each type of environmental signal. The cuticle of mechanosensory bristles is supported by longitudinal pillar-like bulges that enhance the mechanical strength of the bristle so that its deflection caused by mechanical contact or air vibration is sensitively transmitted to the mechanosensory neurons that innervate to the base of the bristle. Cuticles of olfactory bristles contain multiple pores, the nanopore, of 30 to 100 nm in diameter that serves as a molecular filter, allowing the entry of airborne olfactory molecules of up to a few nm and preventing the entry of larger particles of dust and virus (Steinbrecht 1997; Hunger and Steinbrecht 1998; Shanbhag *et al*. 1999). In the case of gustatory sensillum, a single tip pore is used to incorporate water-soluble taste molecules in the food (Shanbhag *et al*. 2001). The corneal nipples are equally spaced ∼200 nm high protrusions covering the corneal lens (Kryuchkov *et al*. 2011). It deflects water droplets, self-cleaning the corneal surface, and decreases light reflection. Despite industrial fabrication mimicking those surface structures attracting much attention from engineers (Bhushan 2009), the investigation into the biological processes of cuticle nanostructure formation and its genetic basis has been slow. One clue for approaching this problem was obtained from the recent study on the *Drosophila* olfactory organ with cuticle nanopores. We previously reported that the gene *Osiris23*/*gore-tex* (*Osi23*/*gox*) is expressed in the olfactory hair cell (trichogen) during day 2 of pupal development when the outermost layer of the epicuticle (envelope) is secreted (Ando *et al*. 2019a). Nanopores were shown to be derived from the indentation of the enveloping layer. In *Osi23*/*gox* mutants, the envelop indentation was flattened, nanopores were lost, and the mutant animals exhibited reduced olfactory response. Since the *Osi23*/*gox* mutant adults are viable and fertile with the normal external shape at the macroscopic level, this gene functions specifically in the nano-level patterning of the cuticles.

*Osi23*/*gox* belongs to the *Osiris* gene family of 25 homologous genes in the *Drosophila* genome. 22 of *Drosophila Osiris* genes are clustered in the chromosome region 83E, corresponding to the triple-lethal region, which shows unusual dosage sensitivity: either one or three copies of the region caused lethality (Dorer et al., 2003; Lindsley et al., 1972). *Osi* gene family was found in many insect genomes spanning the basal groups of mayflies and silverfish to highly derived dipterans. No *Osi* homologs are found in the genomes of basal hexapods (bristletails, Archaeognatha), crustaceans, and other arthropods. Molecular phylogenetic analysis demonstrated that specific classes of *Osi* genes from different insects are clustered. This suggests that the *Osi* gene was acquired in the early stage of insect evolution, rapidly increased in number, and diverged (Shah *et al*. 2012). Then, the gene family was conserved thereafter. This implies that each member of *Osi* genes is conserved in insect evolution. Although a few studies are addressing the function of specific *Osi* genes (Smith *et al*. 2018; Scholl *et al*. 2018, 2023; Scalzotto *et al*. 2022), no comprehensive analysis of the expression and genetic requirement for *Osi* family genes has been reported for *Drosophila* or other insects.

In this study, we performed a gene expression analysis of all *Osi* genes in the *Drosophila* head. The results showed that in the early stage of adult cuticle deposition, *Osi* gene transcripts are found exclusively in specific cuticle-secreting cells in patterns unique for each *Osi* gene. Collectively, most adult cuticles are secreted by cells expressing specific combinations of *Osi* genes. Systematic gene knockout experiments demonstrated a varying degree of requirement, from haploinsufficiency for viability to no apparent requirement of adult viability and fertility. Among adult viable *Osi* mutants, many showed specific defects in olfactory nanopores, gustatory tip pores, and corneal nipples in the eye. Those results indicate that the *Osi* gene family plays an essential role in cuticle nanopatterning in insects.

## 2. Results

### 2.1. Unique combination of *Osiris* gene expression prefigures morphogenesis of specific cuticle structures

Expression patterns of *Osiris* genes in the *Drosophila* embryo were described previously (Ando *et al*. 2019a). We sought to study tissue expression patterns of *Osiris* genes in the head of pupae at 42-44 hours APF (after puparium formation) when the amounts of *Osiris* gene transcripts peaked in the pupal stage (Brown et al., 2014; Larkin et al., 2021; Sobala & Adler, 2016), and the expression of *Osi23*/*gox* was detected in the olfactory hair cells (Ando et al., 2019). This is the time when the envelope layer of the cuticle is assembled before the production of chitin and other components of the procuticle (Ando et al., 2019; Sobala and Adler, 2016). We reasoned that if other *Osiris* genes play a role analogous to *Osi23*/*gox* in nano-patterning of the cuticle through modulation of envelope shape, they are expressed at this stage of envelope formation.

Fluorescence *in situ* hybridization (FISH) was performed on the whole head with 25 probes for each *Osi* RNA (*Osi1* to *Osi24*. *Osi10* was reannotated as *Osi10a* and *Osi10b,* Figure 1A; Supplementary Figure S1; S2). The samples were co-stained with anti-phosphotyrosine and anti-Futsch antibodies to reveal cell outline and bristle shaft cells (trichogen) and neurons, respectively, and the nuclei were labeled with DAPI (Supplementary Figure Si and S2). Based on the low magnification views, *Osiris* expression patterns were classified into three categories (Figure 1A, Supplementary Figure S2, S3, Supplementary Table S1). The first group of genes (*3*, *7*, *9* and *22*) are mainly expressed in epidermal cells, and the second group (*1*, *4*, *5*, *6*, *8*, *11*, *12*, *13*, *16*, *21*, *23* and *24*) are expressed in various sensory organs in the eye, antenna, maxillary palp, and proboscis (Figure 1A). The third group of genes (*2*, *10a*, *10b*, *14*, *15*, *17*, *18*, *19* and *20*; Supplementary Figure S2) were not expressed at a detectable level in the head of this stage. We noted that our FISH assay is sensitive enough to detect robust sensory expression of *Osi16* and *Osi23*/*gox* RNAs that were classified as “low” expressed genes (5 fragments per kilobase of exon per million mapped reads / FPKM) in the modENCODE temporal gene expression database (Graveley et al., 2011). *Osi* expression patterns are highly divergent and complex. We describe cell type-specific expression patterns of 16 *Osiris* genes expressed in the pupal head at 42 hours APF. A detailed expression of each *Osi* gene is presented in Figures 1-4 and Supplementary Figure S3.

**Figure 1.**
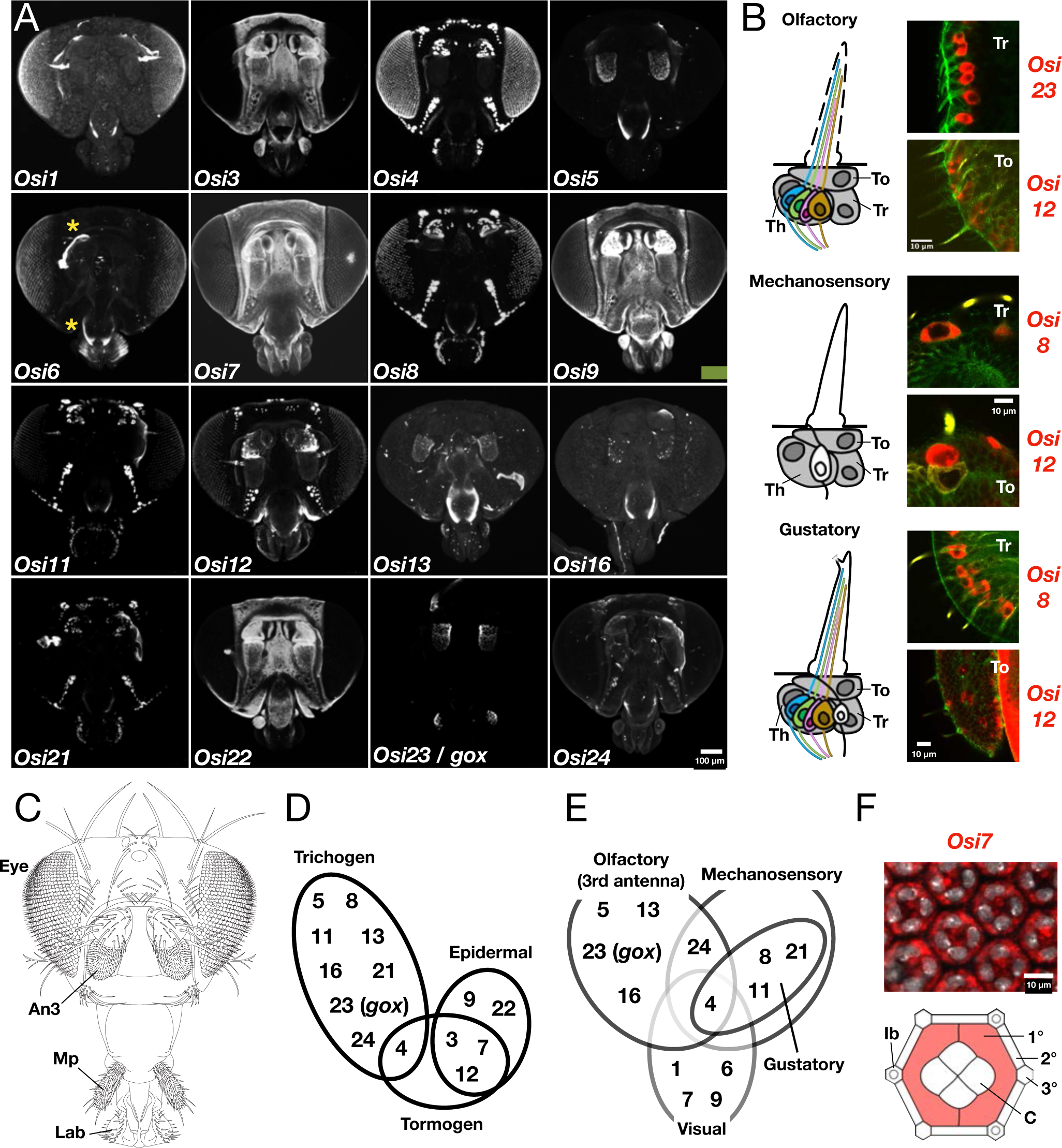
Expression patterns of *Osi* genes in the pupal head. (A) mRNA expression of 16 *Osi* genes in 42 hours APF pupal heads. Asterisk (*) indicates a non-specific signal to the pupal cuticle remnants. (B) Schematics of 3 sensory bristle types and examples of pupal *Osi* gene expressions in To (tormogen cell) and Tr (trichogen cell). Th: thecogen cell. mRNA (red), phosphotyrosin (green, cell junction) and Fusch (yellow, bristle shaft). (C) A schematic of *Drosophila* adult head. An3: third antennal segment. Mp: maxillary palp. Lab: labellum. (D) Summary of *Osi* gene expressions in 3 types of cuticle-secreting cells. (E) Summary of *Osi* gene expressions in different sensory organ types. Note that *Osi* expressions in the gustatory organ are a subset of expressions in the mechanosensory organ. (F) An example of *Osi* expression in the compound eye. *Osi7* mRNAs were detected in primary pigment cells (1°) and cone cells (C). 2°and 3°: secondary and tertiary pigment cell. mRNA (red) and DNA (grey).

### 2.1. Expressions in sensory organs

The head of adult *Drosophila* is highly concentrated with various sense organs (Figure 1B, 1C). The olfactory organs are hair-like protrusions with nanopores covering the surface of the maxillary palp (Mp) and the third antennal segment (An3; Figure 1C, Figure 2G). Taste organs with single-tip pores are present on the surface of the labella (Lab), the bilateral structure in the distiproboscis. 31 gustatory bristles cover the dorsal (outer) side of the labella (Figure 3J). In the ventral side of the labella, rows of taste pegs are present on the side of 6 rows of pseudotrachea, the teeth-like cuticular structures (Figure 3J).

**Figure 2.**
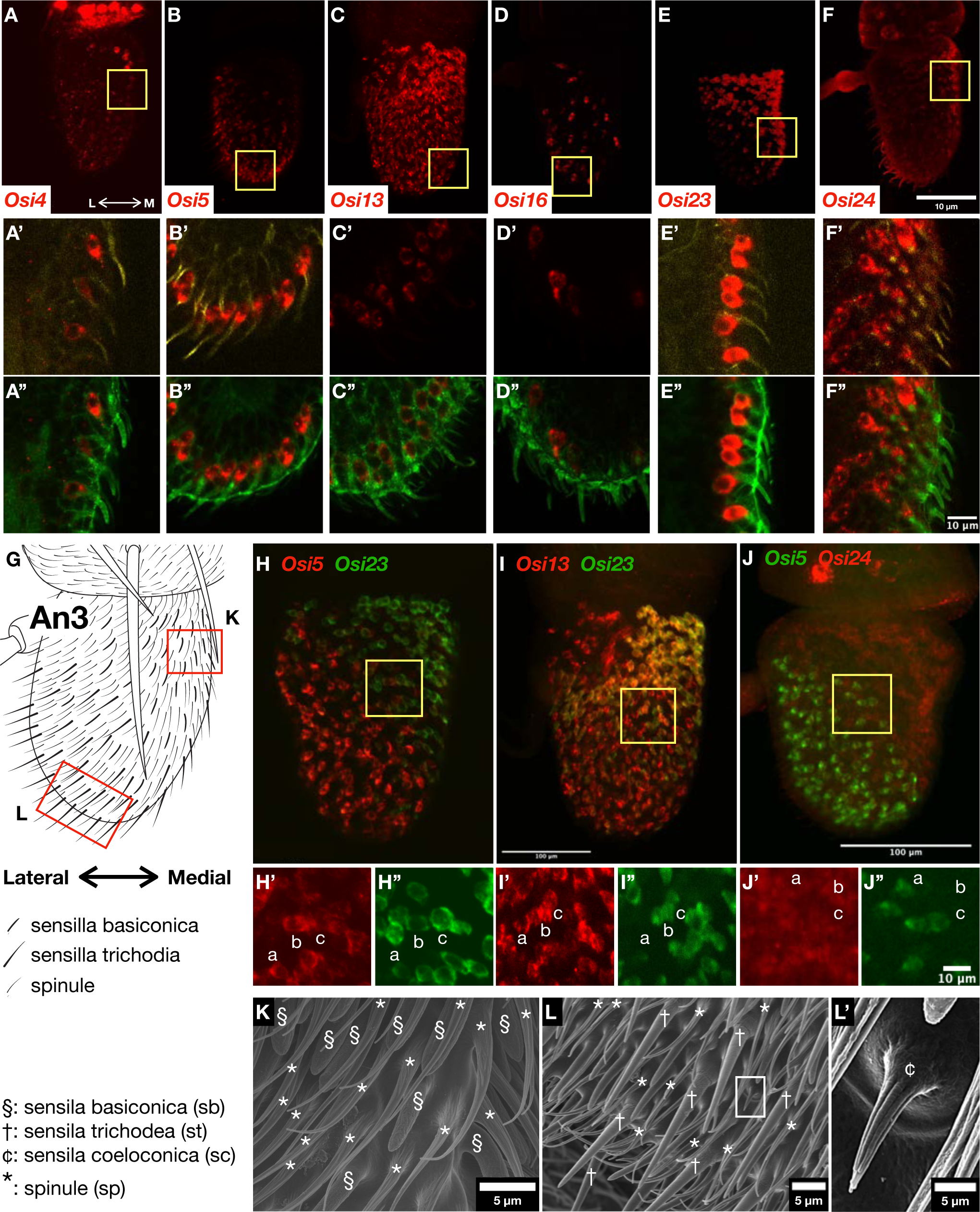
*Osi* gene expressions in olfactory organs. (A-F) Overview of *Osi* gene expressed in the third antennal segment (An3). Lateral (L), medial (M). (A’-F’ and A’’-F’’) Magnified views of yellow boxes in A-F. Phosphotyrosine (green), Futsch (green). (G) A schematic of An3. Sensilla basiconica (sb) and sensilla trichordia (st) are enriched in the medial top and lateral bottom regions, respectively. SEM views of each region are shown in K and L. (H-J) Two-color FISH images of 3 pairs of *Osi* mRNA expression. (H’-J’ and H’’-J’’) High magnification views of the yellow boxes. Note that *Osi23* expression overlaps significantly with *Osi13* (I) but is distinct from that of *Osi5* (H). *Osi24* expression differs from *Osi5* (J). (K) SEM image of the medial-top region enriched with sb. (L) Lateral-bottom region enriched with st. (L’) Enlarged view of sc (white box in L).

**Figure 3.**
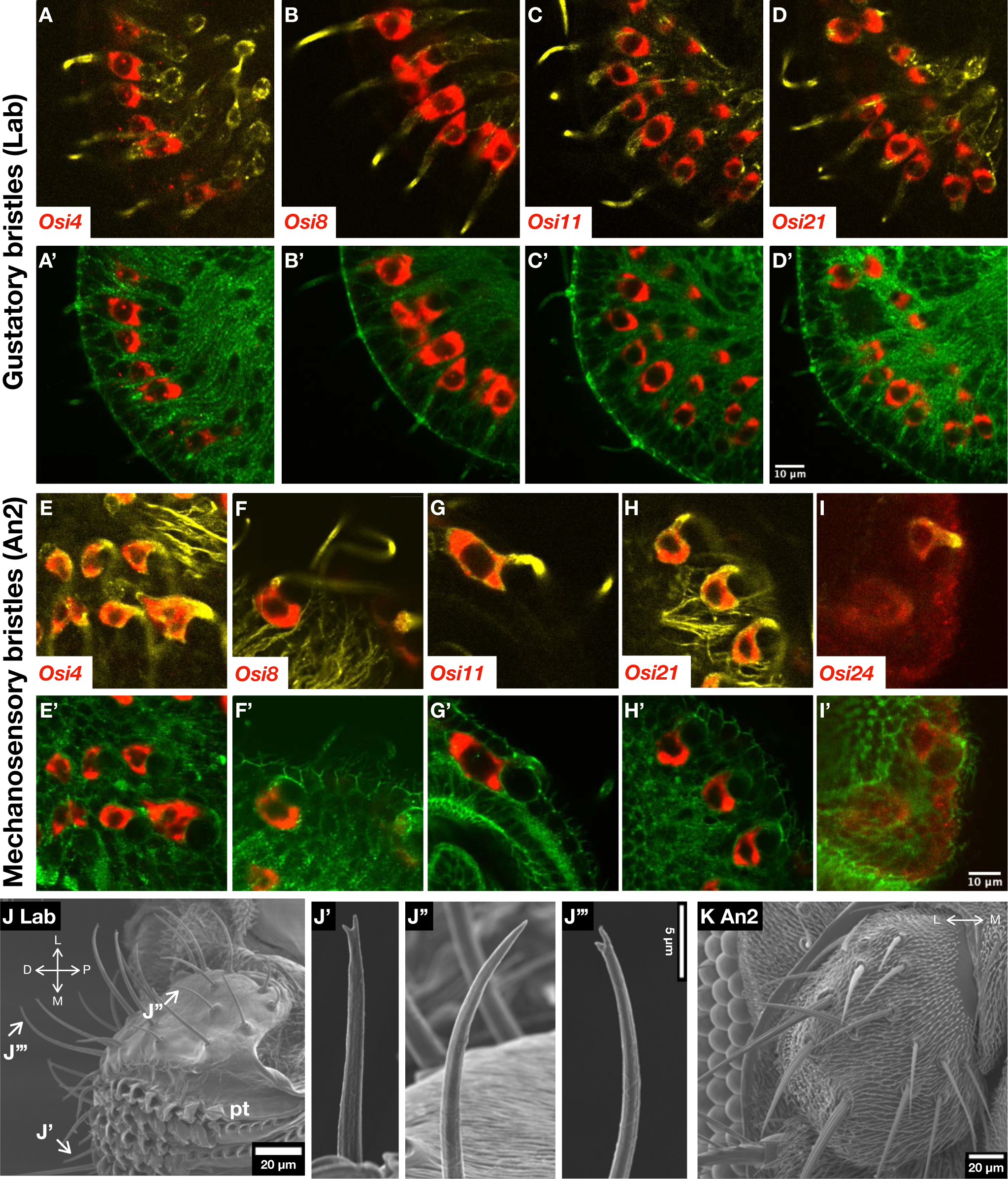
*Osi* gene expressions in gustatory and mechanosensory organs. (A-I) Expressions of *Osi* genes expressed in trichogen co-labelled with Futsch. (A’-I’) *Osi* expressions co-labelled with phosphotyrosine. (J) SEM observation of labellum (right side). J’: small taste bristle. J’’: intermediate taste bristle. J’’’: large taste bristle. Distal (D), proximal (P), pseudotrachea (pt). (K) SEM observation of An2 with mechanosensory bristles.

Mechanosensory bristles of various sizes are present in many positions of the epidermis. External structures of those sensory organs consist of bristle (hair) and socket, secreted by trichogen and tormogen cells, respectively (Figure 3K). The light-sensing compound eye is an assembly of about 800 ommatidia, each covered with a transparent lens cuticle that focuses incoming light to the internal retinal cells (Figure 5G). In addition, three ocelli present on the dorsal head are also covered with lens cuticle.

Nine *Osi* genes are expressed in trichogen cells, which were classified into two partially overlapping groups, one expressed in olfactory organs and the other in mechanosensory organs (Figure 1D, E). The latter group included those expressed in gustatory organs. In addition, four *Osi* genes expressed in lens-secreting cells were grouped into another cluster. The result suggests distinct *Osi* genes are expressed in cells covering sensory organs with different cuticle nanostructures (Figure 1D, E; Supplementary Table S1).

Tormogen cells of mechanosensory, gustatory, and olfactory organs expressed *Osi3*, *Osi4*, *Osi7*, and *Osi12*, which form a group overlapping the epidermis- and trichogen-expressed genes (Figure 1C). *Osi3* and *Osi7* are also expressed in the epidermis. The result suggests that *Osi* genes do not distinguish socket cuticles of different sensory organs.

### 2.2 Olfactory sensillum

Olfactory organs are categorized into sensilla basiconica (sb), sensilla trichordia (st) and sensilla coeloconica (sc), each has multiple cuticular nanopores (Shanbhag et al., 1999, Figure 2K, L, L’). Those sensilla are further classified based on size, expression of olfactory receptors, and response to specific chemicals (Chai *et al*. 2019). All three types of olfactory sensillum are present in An3, and only sb is present in Mp. *Osi13*, *Osi23*/*gox*, and *Osi24* were detected in trichogen cells of maxillary palp in patterns resembling the distribution of HA-Gox driven by the *Osi23*/*gox* promoter (Figure 1B; Figure 2; Supplementary Figure S3; Ando et al., 2019a), suggesting that those genes are expressed in sb. *Osi5* was expressed abundantly in An3, in a pattern complementary to the sb location labeled by strong *Osi23* (Figure 2H). Based on the similarity to the st distribution in adult an3, *Osi5* is likely to be expressed in st. Osi13 and Osi23 were expressed in mostly overlapping patterns in An3 (Figure 2I) and Mp (Supplementary Figure S4). Osi24 expression partially overlapped with Osi5 (Figure 2J). *Osi16* was expressed in a scattered pattern in An3 (Figure 2D; Supplementary Figure S3). It is possible that *Osi16* is expressed in sc. Each olfactory sensillum is associated with sets of neurons expressing specific olfactory receptors or Ionotropic receptors (reviewed by Vosshall and Stocker, 2007). However, none of those receptor promoters of Gal4 fusion are active in ∼42 hours APF. Since no specific marker for sb, st and sc expressed in this stage is available, definitive identification of *Osi5* and *Osi16* expressing cells requires new lineage tracing tools.

### 2.3 Mechanosensory and gustatory sensillum

Mechanosensory bristles transmit mechanical stimuli to the mechanoresponsive nerve terminus attached to one side of the bristle base. Their shape is characterized by prominent bulges running along the long axis of the bristle, which is pre-patterned by actin bundles formed during the pupal stage (Figure 3K, Lees AD and Picken L. E. R., 1945; Tilney et al., 1995). Gustatory organs sense water-soluble chemicals with taste neurons inside of the bristle. Long and short types of gustatory bristles are branched, and one of the branches has a pore at the tip, through which water and dissolved taste molecules reach gustatory neurons inside the bristle (Figure 1B, Figure 3J’. Gustatory bristles are also innervated by the mechanosensitive neuron (Jeong et al., 2016). Five *Osiris* genes (*Osi4*, *Osi8*, *Osi11*, Osi21 and *24*) are expressed in all trichogen cells of the mechanosensory bristles (Figure 1B; Figure 3; Supplementary Figure S1). Four of these (*Osi4*, *Osi8*, *Osi11* and *Osi21*) were detected in all trichogen cells of gustatory bristles of the proboscis (Figure 1B, 1E; Figure 3; Supplementary Figure S3). In addition, *Osi3* expression was detected in gustatory tormogen cells (Supplementary Figure S3).

### 2.4 *Osi* expression in the eye

*Osi4*, *Osi6*, *Osi7* and *Osi9* are expressed in primary pigment cells of the compound eye (Figure 1F; Figure 4; Supplementary Figure S3). *Osi7* was also expressed in cone cells (Figure 4). Those cells are involved in the secretion of the transparent lens cuticle. In addition, *Osi4* expression was detected in unidentified cells deep in the compound eye (Supplementary Figure S3). It is unlikely that those cells are involved in lens secretion.

**Figure 4.**
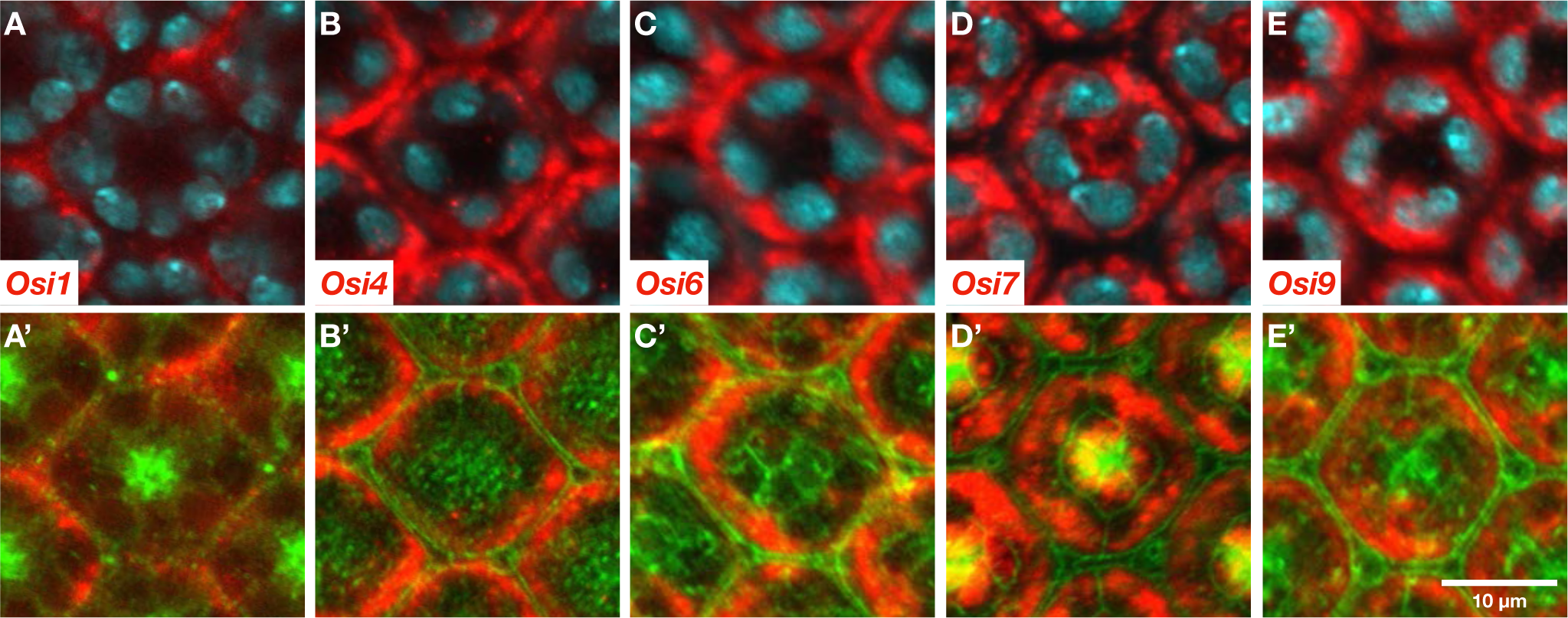
*Osi* gene expressions in the compound eye. (A-E) *Osi* expressions with nuclei. (A’-E’) *Osi* expressions with phosphotyrosine. Approximate depth from the apical surface: 4.86 μm (*Osi1*), 1.62 μm (*Osi4*), 1.62 μm (*Osi6*), 2.16 μm (*Osi7*), 1.62 μm (*Osi9*).

### 2.5 Epidermis and arista

*Osi3*, *Osi7*, *Osi9* and *Osi22* are expressed in most parts of the epidermis (Figure 1A), which is covered by an epidermal protrusion called spinule (sometimes called trichome or hair). We noted that a part of the central-posterior part of the proboscis showed little or no expression of any of the *Osi* genes (Supplementary Figure S3). Another position lacking *Osi* expression was a horizontal strip above the antenna (Figure 1C). This region appears to be located dorsal to the ptilinum and folded inside the adult head.

*Osi1*, *Osi8*, *Osi11* and *Osi12* are expressed in arista, the distal part of the antenna (Supplementary Figure S3). *Osi1*, *Osi8* and *Osi11* are expressed in the basal cylinder and farther distal part, with enhanced expression in the dorsal side. *Osi12* showed a distinct pattern of a ring in the boundary of the basal cylinder and arista.

### 2.6 Mutagenesis of *Osiris* gene family

Mutations of a subset of *Osiris* genes have been reported (Ando *et al*. 2019a; Scalzotto *et al*. 2022; Scholl *et al*. 2023), but no comprehensive mutagenesis of the *Osiris* gene has been performed before. Preliminary experiments to knock down *Osi* genes with transgenic UAS-RNAi strains (Dietzl *et al*. 2007; Ni *et al*. 2008) gave mixed results: some RNAi constructs caused lethality while others did not (Supplementary Table S2). We then performed systematic gene knocked out of all 25 *Osiris* genes using the transgenic guide RNA and Cas9 technique (Kondo and Ueda, 2013; Table 1, Supplementary Table S3). Multiple small deletion alleles causing frameshift mutations in the open reading frame of each *Osiris* were recovered for 24 *Osiris* genes, among which 5 were lethal (*Osi7*, Osi*17*, *Osi20* and *Osi24*) and two semi-lethal (*Osi10a, Osi14*). We were unable to recover any mutation of *Osi6* with 3 different guide RNA design. One allele of lethal *Osi6* previously isolated was embryonic lethal with a strong cuticle defect (Ando *et al*. 2019a). Heterozygous *Osi6* and *Osi7* stocks were weak and sluggish, indicating that one dose reduction of those genes seriously impacted viability.

**Table 1.** List of *Osiris* gene mutants.

| Symbol | Viability | Viable allele* | adult phenotype | Comment |
| --- | --- | --- | --- | --- |
| *Osi1* | V | 9/10 |  |  |
| *Osi2* | V | 2/2 |  |  |
| *Osi3* | V | 7/7 |  |  |
| *Osi4* | V | 4/4 | compound eye |  |
| *Osi5* | V | 5/7 |  |  |
| *Osi6* | L | No allele recovered. |  | Lethal (Ando et.al. 2019) |
| *Osi7* | L | 0/3 |  | Lethal (Ando et.al. 2019) |
| *Osi8* | V | 5/5 |  |  |
| *Osi9* | V | 6/6 | compound eye | Viable (Scholl et al., 2023) |
| *Osi10a* | semi-L** | 3/4 |  |  |
| *Osi10b* | V | 4/5 |  |  |
| *Osi11* | V | 4/4 | gustatory organ |  |
| *Osi12* | V | 4/6 |  |  |
| *Osi13* | V | 3/5 |  |  |
| *Osi14* | semi-L** | 4/5 |  |  |
| *Osi15* | V | 3/3 |  | Viable (Scholl et al., 2023) |
| *Osi16* | V | 5/5 |  |  |
| *Osi17* | L | 0/7 |  |  |
| *Osi18* | V | 7/7 |  |  |
| *Osi19* | V | 6/7 |  | Semi-lethal (Ando et.al., 2019; Scholl et al., 2023) |
| *Osi20* | L | 0/3 |  |  |
| *Osi21* | V | 2/2 |  | Suppressed norpA phenotype. Lee et al., (2013) |
| *Osi22* | V | 4/6 |  |  |
| *Osi23* | V | 3/3 | Olfactory organ (sb, st). | Viable (Ando et.al. 2019) |
| *Osi24* | L | 0/6 |  |  |

\* Count of viable allele among total number of alleles with out of frame in/del mutation.
\*\*Frequent dead pupae were observed.

Heads of adult viable mutant animals were examined by field emission scanning electron microscopy (FE-SEM). Cuticle patterns of the antenna, compound eyes, and proboscis were observed at magnifications up to 10,000x (Figure 5). External morphology was observed in the antenna (olfactory organs in An3, mechanosensory organs in An2, arista), Mp, Lab (gustatory organs, pseudo trachea), and the lens of compound eyes of multiple independent alleles of homozygous mutant adults of each gene (Table 1, Supplementary Table S3). Defects in nanostructures were found in olfactory organs, gustatory organs, and eye lenses, as described below.

**Figure 5.**
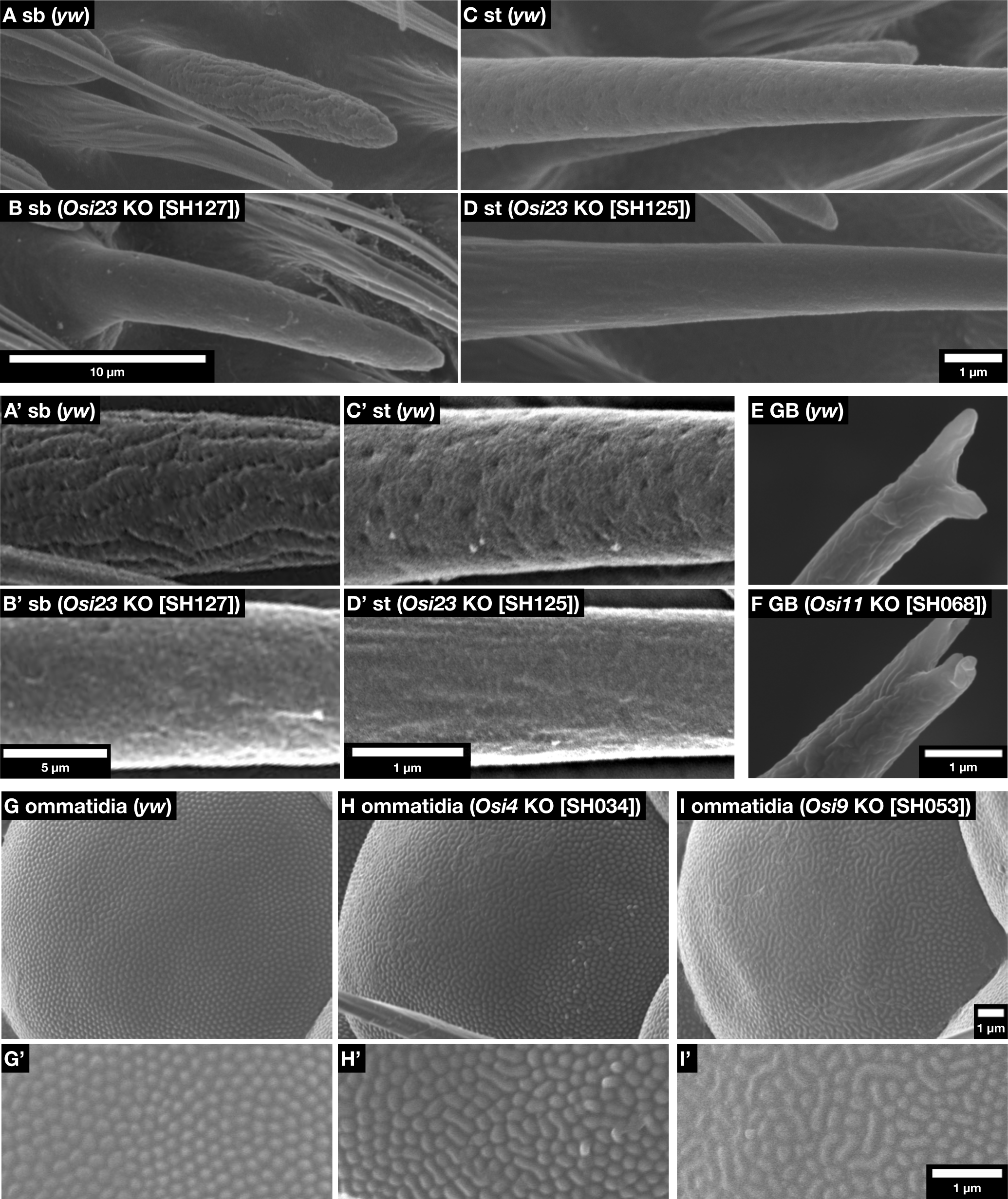
Impact of *Osi* gene mutations on cuticle nanostructure formation. (A, A’, B, B’) SEM images of sb in An3. (C, C’, D, D’) st in An3. Note the clear loss of nanopores in sb and st. (E, F) Tip of long gustatory hairs. In *Osi11* KO, each tip is further bifurcated. (G-I, G’-I’) Surface views of ommatidium. Note that individually separated nipple arrays in control (G) are laterally fused in *Osi4* and *Osi9* mutants.

### 2.7 Phenotypes in the olfactory organs

*Osi23*/*gox* mutants show the loss of nanopores in the sb of maxillary palp, as previously reported (Ando *et al*. 2019a). The mutations also caused the loss-of-nanopore phenotype in the sb of An3 (Figure 5). Furthermore, we observed a loss of nanopores in the st of An3 (Figure 5). We also examined olfactory organ phenotypes in mutants *Osi4*, *Osi5*, *Osi13*, *Osi16* and *Osi24* expressed in bristles of olfactory organs in the An3, but no obvious phenotype was observed. To investigate the role of *Osi23*/*gox* in st morphogenesis, we re-examined its expression pattern in An3 at 42 hours APF and found that many trichogen cells express *Osi23*/*gox* RNA at a low level in the ventrolateral region of An3, which is covered by numerous st in the adult (Figure 6). The results imply that *Osi23*/*gox* contributes to nanopore formation in the two types of olfactory hair cells of sb and st. The external appearance of sc (sensilla coeloconica) was normal in *Osi23*/*gox* mutants.

**Figure 6.**
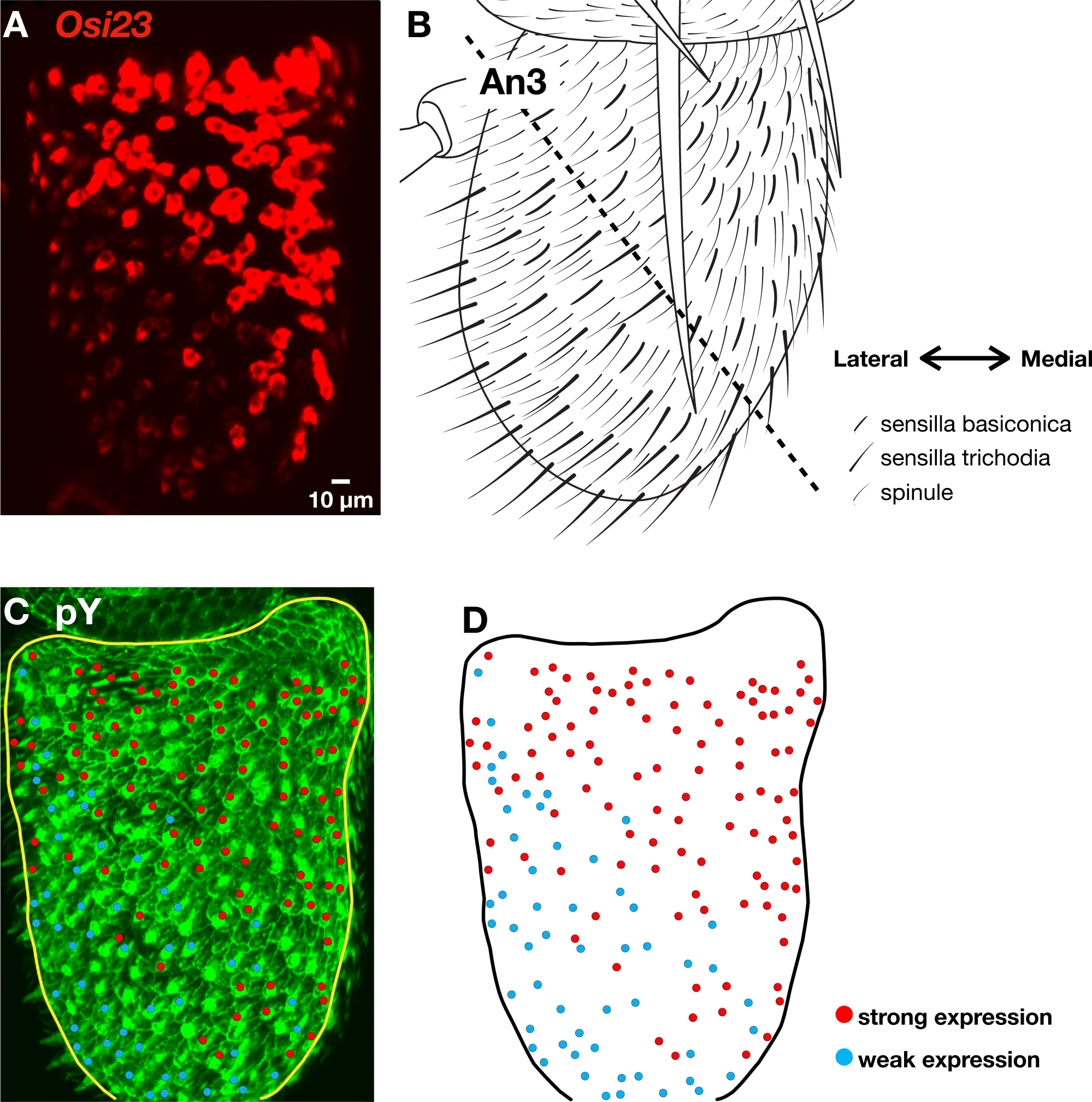
*Osi23* expression in An3. Cut-open views of the top surface of An3. (A) *Osi23* is strongly expressed in the top-medial area enriched with sb and weakly in the bottom-lateral area enriched with st. (B) Schematics of An3. (C, D) Anti-phosphotyrosine (pY) staining and the map of *Osi23* expression.

### 2.8 Phenotypes in the gustatory organ

Among 4 *Osi* genes expressed in the gustatory organs (*Osi4*, *Osi8*, *Osi11* and *Osi21)*, mutants of *Osi11* showed a change in the morphology of branched tips of long-type and short-type gustatory hairs (Figure 5).

### 2.9 Phenotypes in the lens

In control eyes, corneal nipples are ∼30 nm high protrusions spaced by ∼255 nm equally spaced on the surface of the lens (Kryuchkov *et al*. 2011). In *Osi4* and *Osi9* mutants, some corneal nipples are fused laterally to form a labyrinthine pattern. No specific defect was observed in *Osi6* and *Osi7*.

### 2.10 Phenotype of *Osi17* knockdown in the wing

Although *Osi17* was homozygous lethal, RNAi-mediated knockdown by the *actin-Gal4* driver caused an eclosion defect with shrunken wings (Supplementary Figure S5). Since Osi17 is expressed in the embryonic tracheal system (Ando et al., 2019), we targeted RNAi to the tracheal system using the tracheal driver. We did not observe the wing defect. We next selectively produced *Osi17* mutant clones using *Ubx*-*flip* recombinase. The mutant flies reproduced the shrunken wing phenotype. Since recombination occurred in the wing pouch region but not in the trachea or adult muscle precursor cells associated with the wing disc, we conclude that the *Osi17* function is required in the wing epithelium to produce a properly expanded wing.

## 3. Discussion

In this study, we presented the expression patterns of *Osiris* gene mRNAs in the pupal heads at the earliest stage of cuticle formation. Of 25 *Osi* genes, 16 were expressed in specific patterns in cuticle-secreting epidermal and sensory organ cells. Those cells form fine protrusions (trichomes and spinules of epidermis), corneal nipples (eye lens), ridges (mechanosensory bristles), tip pores (gustatory bristles), and nanopores (olfactory bristles). Some parts of the epidermis, such as the rear part of the proboscis, lack cuticular protrusions. No *Osi* gene expression was observed in this part. The results imply that the *Osi* gene family is a strong candidate for the regulator of various forms of cuticle nano-patterns.

### 3.1. *Osiris* functions in sensory bristle nanopatterns

Nine *Osi* genes are expressed in sensory bristle-forming trichogen cells. Two showed clear defects in cuticle nano-patterns. In the olfactory bristles in An3 and Mp, *Osi23*/*gox* was required not only for the nanopore formation of sensilla basiconica (Ando et al., 2019a; this study) but also for sensilla trichordia. This is consistent with the expression of *Osi23*/*gox* in bristle cells, albite at a low level, in the ventrolateral region of An3, where st is enriched (Figure 6). Although *Osi5* is likely to be expressed in st, *Osi5* mutants did not show obvious defects in nanopores of st or other olfactory bristle types, so as the other viable mutants of *Osi* genes expressed in olfactory bristle hair cells in An3 and Mp (*Osi4*, *Osi13* and Osi*16*).

Of 4 *Osi* genes expressed in gustatory and mechanosensory bristle cells (*Osi4*, *Osi8*, *Osi11* and *Osi21*), only *Osi11* mutants showed defects in the formation of the forked bristle tip morphology in the gustatory bristles. No change in mechanosensory bristles shape was observed. The results support the hypothesis that the *Osi* family genes play important roles in cuticle nanopatterns of sensory bristles. The results also showed that the other trichogen-expressed *Osi* genes are apparently dispensable for cuticle patterning when singly mutated. It is possible that some *Osi* genes function redundantly in those cells, as demonstrated for the embryonic trachea where mutations of three genes (*Osi9*, *Osi15* and *Osi19*) were required to produce clear tracheal phenotype (Scholl et al., 2018).

*Osi3*, *Osi4* and *Osi12* are expressed in socket-forming tormogen cells, but none of their mutants showed visible defects in the socket.

### 3.2. *Osiris* functions in the corneal nipple nanopattern

Of the 5 *Osi* genes expressed in the lens cuticle-secreting cells, mutants of *Osi4* and *Osi9* showed defects in the pattern of nipple arrays. Lateral fusion of nipple arrays formed labyrinthine patterns that are reminiscent of the lens patterns observed in some *Drosophila* species and in *Drosophila* melanogaster mutants deficient in Retinin and waxes that partly constitute the nipple structures (Blagodatski *et al*. 2015; Kryuchkov *et al*. 2020). It is likely that *Osi4* and *Osi9* are components of the reaction-diffusion mechanism of corneal nipple array patterning (Kryuchkov et al., 2020; Turing, 1990).

### 3.3. *Osiris* gene functions in the epidermis

*Osi3*, *Osi7*, *Osi9* and *Osi22* are expressed strongly in the epidermis. Although three of them (*Osi3*, *Osi9* and *Osi22*) are viable and did not cause obvious defects in the epidermal cuticle and trichome, it was previously shown that embryonic lethal *Osi6* and *Osi7* mutants showed strong defects in the larval cuticle formation (Ando *et al*. 2019a). In addition, the lack of *Osi17* function caused wing expansion defects, likely due to the weakening of the epidermal cuticle. Whether those defects reflect the functions of cuticle nanopatterns or general cuticle production remains to be determined.

### 3.4. Dynamic *Osiris* gene expression

In the case of *Osi4*, we observed related but distinct expression patterns in a batch of pupae similarly staged, fixed at 42 hours APF, and processed together in a single tube (Supplementary Figure S3). One head shows a high expression in the eye but not in the pseudotrachea. Another one showed weaker eye expression and prominent expression in pseudotrachea. It is possible that *Osi4* expression dynamically changes, and a slight difference in the developmental stage (less than +/-0.5 hours) causes a significant difference in the expression pattern. As described above, we noted that nanopore formation in sensilla trichordia (st) is sensitive to *Osi23*/*gox* mutation, although its expression in st was low at 42 hours APF. A stage of high *Osi23*/*gox* expression is st primordia may have been missed. Time-course analyses of the expression *Osi23*/*gox* and other *Osi* genes are required to fully understand how the genes contribute to the complex morphogenesis of cuticle nano-patterns.

### 3.5. Genetic requirement for *Osi* genes

Systematic knockout of *Osi* genes revealed variable requirements for each *Osi* gene in organismal viability and cuticle nano-patterning. The requirement for *Osi6* and *Osi7* activities is especially high since heterozygosity of either of the genes reduces the fitness of animals (Ando et al., 2019a; this study). Five additional *Osi* genes are lethal or semi-lethal. The results support the hypothesis that the combined effect of *Osi* genes accounts for the haplo-insufficiency of the chromosomal locus 83D-E covering the complex of 22 Osi gene (Lindsley et al., 1972; Shah et al., 2012).

The 22 *Osi* genes are densely packed and sometimes overlap in the ∼168 kb region of 83D-E. In such cases, enhancer sharing, and co-regulation would be anticipated. However, we did not see any obvious co-regulation of neighboring genes. 5 kb *Osi23/gox* genomic fragment with 2kb each of 5’ and 3’ fragment flanking the gene was sufficient to rescue *Osi23/gox* mutant phenotype (Ando et. al., 2019). Therefore, the expression of each *Osi* gene is likely to be regulated independently by nearby enhancer sequences.

Although the mutations in *Osi4*, *Osi9*, *Osi11* and *Osi23*/*gox* caused defects in cuticle nano-patterns in the compound eye, and gustatory and olfactory bristles, respectively, mutations of other *Osi* genes co-expressed in those tissues did not cause a notable change in the cuticle. No defect in the cuticle pattern of mechanosensory organs and epidermis was observed, although expression of multiple *Osi* genes was detected. Genetic redundancy, reported for the tracheal function (Scholl *et al*. 2023), is a likely reason. The expression patterns reported in this work will guide a future study of introducing multiple mutations in genes co-expressed in the same cell type.

The laterally fused corneal nipple phenotypes of *Osi4* and *Osi9* are reminiscent of the eyes of some *Drosophila* species (other than *D. melanogaster*) that show naturally fused nipple patterns with decreased surface wetting and increased light reflection compared to *D. melanogaster* (Kryuchkov *et al*. 2020). It would be interesting to study the difference in the functions of *Osi4* and *Osi9*-related genes in those *Drosophila* species to investigate the potential function of *Osi* genes in the variation of cuticle nanopatterns.

## 5. Materials and Methods

### 5.1. Key resource table

| REAGENT or RESOURCE | SOURCE | IDENTIFIER |
| --- | --- | --- |
| Antibodies |  |  |
| P-Tyr-1000 MultiMab Rabbit mAb mix | Cell Signaling Technology | Cat# 8954, RRID:AB_2687925 |
| futsch antibody | DSHB | Cat# 22C10, RRID:AB_528403 |
| Goat anti-Rabbit IgG, Alexa Fluor 488 | Thermo Fisher Scientific | Cat# A-11034, RRID:AB_2576217 |
| Goat anti-Mouse IgG, Alexa Fluor 633 | Thermo Fisher Scientific | Cat# A-21052, RRID:AB_2535719 |
| Anti-Digoxigenin-POD, Fab fragments | Roche | 11207733910 |
| Streptavidin-POD | Roche | 11089153001 |

| Chemicals, Peptides, and Recombinant Proteins |  |  |
| --- | --- | --- |
| Alexa Fluor 568 Phalloidin | Thermo Fisher Scientific | Cat# A12380 |
| TSA Fluorescein | PerkinElmer | SAT701001KT |
| TSA Cyanine 3 | PerkinElmer | SAT704A001KT |
| Paraformaldehyde | TAAB | Cat# P001 |
| Glutaraldehyde EM Grade | TAAB | Cat# G011/1 |
| 16% Formaldehyde Solution (w/v) | Thermo Scientific | Cat# 28908 |
| Blocking Reagent | Roche | Cat# 11 096 176 001 |
| DIG RNA Labeling Mix, 10x conc. | Roche | Cat. No. 11 277 073 910 |
| Biotin RNA Labeling Mix, 10x conc. | Roche | Cat. No. 11 685 597 910 |
| Antifade Mounting Medium with DAPI | Vector Laboratories | H-1200 |
| Xylene | Wako | 244-00086 |
| Acetone | Wako | 016-00346 |
| Ethanol | Wako | 057-00451 |
| PBS (10×) | nacalai tesque | Cat# 27575-31 |
| Molecular Sieves 3A 1 / 16 | nacalai tesque | Cat# 04170-15 |
| Experimental Models: Organisms/Strains |  |  |
| *D. melanogaster: Oregon R* |  | N/A |
| *D. melanogaster: neur-Gal4* | (Bellaïche *et al.* 2001) | FBti0017282 |
| *D. melanogaster: act-Gal4* | Direct fusion of Gal4 to Act5C promoter | N/A |
| *D. melanogaster: da-Gal4* | (Wodarz *et al.* 1995) |  |
| *D. melanogaster: btl-Gal4* | (Shiga *et al.* 1996) | DGRC 109128 |
| *D. melanogaster: y[1] w{\*}* | Akira Nakamura |  |
| *D. melanogaster: y[1] w[\*]*<br>*P{w[+mC]=Ubx-FLP}1* | Jurgen Knoblich | RRID:BDSC_42718 |
| *D. melanogaster: w[1118];*<br>*P{ry[+t7.2]=neoFRT}82B P{w[+mC]=Ubi-*<br>*GFP(S65T)nls}3R*<br>*P{ry[+t7.2]=A92}RpS3[Plac92]/TM6C,*<br>*Sb[1]* | Bloomington *Drosophila* Stock Center | RRID:BDSC_5627 |
| *D. melanogaster: RNAi strains* | See supplementary Table S1 |  |
| *D. melanogaster: Osi knockout strains* | See supplementary Table S1 | This study |
| Microscope |  |  |
| Confocal microscope | Olympus | FV1000 |
| FE-SEM | JEOL | JSM-IT700HR |
| Software and Algorithms |  |  |
| ImageJ-Fiji | LOCI | https://fiji.sc |

### 5.2. Experimental Models

All *Drosophila* strains were cultured in standard yeast-cornmeal media at 25°C. Fly pupae at the white prepupal stage were picked up and staged.

### 5.3. RNA probe preparation

Antisense RNA probes for each *Osiris* gene were amplified from DNA templates PCR-amplified from the genomic DNA of *y w* strain or *Osiris* cDNA clones. Digoxygenin or biotin-labelled probes were synthesized using the labelling kit (Roche). The template DNA for the *Osi10b* RNA antisense probe was amplified with the primer set (Forward: GTGGCGCGTCGTTTTACTAC, Reverse: TAATACGACTCACTATAGGGCTTGATCGAGGCCCAGCTC). Primer sequences for other genes and the probe preparation method were previously described (Ando et al., 2019).

### 5.4. Fixation of *Drosophila* pupa for FISH

The pupal heads for the FISH experiment were prepared from pupae at 42 hours APF. Pupae were removed from the pupal case and were poked at the posterior abdomen to increase the permeability of the fixative. Pupae were transferred into ∼250 μl of 4% paraformaldehyde in PBS and incubated overnight at 4°C. The pupal cuticle was then removed with fine forceps. Pupal heads (with legs and wings) were collected and rinsed with PBST (0.1% Tween-20 in 1x PBS). Fixed pupal heads were dehydrated by washing for more than 10 minutes with 25%, 50%, 80% and 100% ethanol. After one more wash with 100% ethanol, they were stored at -20°C. Unless otherwise indicated, the incubations and rinses performed below were all at room temperature with 500 μl of each solution. All rinse steps of at least 5 minutes were performed.

### 5.5. Single-color FISH

The procedure modifies the protocol previously described (Inagaki *et al*. 2005). Fixed and dehydrated pupal heads were incubated in a 2 ml microfuge tube with a 1:1 mixture of xylene and ethanol for 60 minutes. Heads were rinsed twice in 100% ethanol and rehydrated through a graded series of ethanol: 80%, 50%, 25% ethanol, and water. Rehydrated pupal heads were incubated in a 4:1 acetone-water solution at -20°C for 10 minutes. Subsequently, heads were rinsed twice with PBST and re-fixed in 4% paraformaldehyde in PBS for 20 minutes. Then rinsed in PBST five times. The pupal heads were prehybridized with prehybridization solution (50% formamide, 5X SSC, 100 μg/ml heparin, 0.1% Tween-20, 100 μg/ml yeast RNA, 10 mM DTT) at 61.7°C for 60 minutes. The prehybridization solution was replaced with the hybridization solution (50% formamide, 5X SSC, 100 μg/ml heparin, 0.1% Tween-20, 100 μg/ml yeast RNA, 10 mM DTT, 10% dextran sulfate) with a final concentration of 0.6 ng/μl digoxigenin-labeled *Osiris* RNA probe. Heads were hybridized overnight at 61.7 °C in a rocking incubator. The pupal heads were washed in a series of wash solutions (50% formamide in PBST), mixed with 5x, 4x, 3x, 2x, and 1xSSC. Each wash was repeated three times for 5 minutes at 61.7°C. Then, heads were rinsed for 5 minutes in PBST five times and incubated in Blocking Reagent (Roche, 1:5000 dilution) for 60 minutes. Then, the heads were incubated in the mixture of Anti-Digoxigenin-POD (Roche, 1:500 dilution), anti-Futsch (DSHB, 1:10 dilution), and Phospho-Tyrosine (Cell Signalling Technology, 1:200 dilution) in PBST for overnight at 4°C. The heads were rinsed five times in PBST at room temperature and then incubated in 50x diluted Cy3 Tyramide Reagent (PerkinElmer Life Science, Inc. 1:50 dilution) in Amplification Dilution buffer for 90 minutes at room temperature. The reaction was terminated by rinsing in Blocking Reagent three times, 10 minutes each. Subsequently, the samples were incubated for 90 minutes in anti-Rabbit Alexa Fluor 488 (Invitrogen, 1:500 dilution) to detect Phospho-Tyrosine and anti-Mouse Alexa Fluor 633 (Invitrogen, 1:500 dilution) to detect Futsch. Finally, the samples were rinsed in PBST three times and mounted in Antifade Mounting Medium with DAPI (VECTASHIELD).

### 5.6. Two-color RNA FISH

After prehybridization and blocking, pupal heads were incubated overnight in the hybridization solution with 0.6 ng/μl each digoxigenin- and biotin-labelled RNA probe. After washing and blocking, the samples were incubated overnight with Streptavidin-POD conjugate (Roche, 1:500 dilution) and then washed. A Tyramide amplification reaction (Cy3) was performed (see in single-color fish). After the reaction, the sample was treated with 0.01 M HCl for 10 minutes to inactivate HRP (Lécuyer *et al*. 2008). The samples were rinsed with PBST, and the blocking was repeated. Then, anti-digoxigenin-POD and anti-Futsch (or phospho-tyrosine) were added simultaneously and incubated overnight at 4°C. The immune reaction was ended by rinsing in PBST three times. Another Tyramide amplification reaction (FITC) was performed for 90 minutes (PerkinElmer Life Science, Inc. 1:50 diluted in Amplification Dilution buffer). After the TSA reaction, the samples were washed and incubated in 1:500 anti-Mouse Alexa Fluor 633 to detect Futsch (or 1:500 anti-Rabbit Alexa Fluor 633 to detect Phospho-Tyrosine) for 90 minutes. Finally, the samples were washed with PBST three times and mounted by Antifade Mounting Medium with DAPI.

### 5.7. Imaginal disc staining

Third instar larvae of the *Osi17* mutant mosaic experiment were dissected, and the wing, haltere, and hind leg discs were fixed at 4% PFA in PBS for 40 min at room temperature. The tissues were blocked with 0.1% BSA in PBST (0.1% Triton-X in PBS) three times for 10 minutes each. Discs were incubated with 1:400 diluted Alexa Fluor 568 Phalloidin in PBST with BSA for 1 hour. Finally, the discs were washed three times and mounted using an Antifade Mounting Medium with DAPI.

### 5.8. Sample preparation for FE-SEM

The adult heads were dissected in PBS and then rinsed with 0.1 M cacodylate buffer 3 times, more than 5 minutes each, and incubated in fixation buffer 1 (2% paraformaldehyde, 2.5% glutaraldehyde, 0.1 M cacodylate buffer) at 4°C overnight. The samples were rinsed with 0.1 M cacodylate buffer 3 times at room temperature, more than 5 minutes each. Then, the samples were incubated in fixation buffer 2 (1% osmium tetroxide, 0.1 M cacodylate buffer) on ice for 120 minutes in a light-shielded condition. The fly heads were further rinsed in water three times on ice with the light-shielded condition and subsequently dehydrated in a gradient of ethanol concentration, from 25%, 50%, 75%, 80%, 90%, 95%, 99.5%, and 100% for 10 minutes each at room temperature. The final 100% ethanol was dehydrated by the addition of a molecular sieve. The samples were dried by overnight incubation in a vacuum. After dehydration, the heads were mounted on double-sided carbon tape on a brass pedestal and coated with OsO4 at approximately 13 nm thickness using an osmium coater (Tennant 20, Meiwafosis Co., Ltd.).

### 5.9. Image acquisition

Fluorescent images were captured via a confocal microscope (Olympus, FV1000) with a 10X objective lens (NA 0.40) for whole pupal head scans and a 60X water immersion objective lens (NA 1.20) for higher resolution images of the antenna, the palps, the distiproboscis and the eyes. For higher resolution images, 0.54μm z stacks were taken. All image data were analyzed via Fiji ImageJ.

External views of adult flies were observed using a field emission scanning electron microscope (JSM-IT700HR, JEOL). A Helium Ion Microscope (ORION Plus, Carl Zeiss, installed at the Nano-processing facility in AIST Tsukuba, Japan) was used in the early screening stage.

### 5.10. Image processing

In order to map *Osi23*/*gox* expression in the curved surface of An3, we employed the ImageJ plugin “SheetMeshProjection” (Wada and Hayashi 2020). This tool allows the conversion of the curved surfaces of objects into 3D stacks of cut-open flat views. The correlation of olfactory organs identified by the phosphotyrosine staining and the strong and weak *Osi23*/*gox* expression was confirmed by moving through the stacks.

### 5.11. Genome editing

Gene knockout strains were produced by the transgenic guide RNA and Cas9 method (Kondo and Ueda, 2013). Multiple alleles were recovered for each gene. Complementation tests with a deficiency chromosome were not possible due to haploinsufficiency of the locus (Lindsley *et al*. 1972). Judgment of lethality was made when all alleles were homozygous lethal (Table 1, Supplementary Table S3). Knock-out strains of *Osi6* were not recovered after the trial with two different guide RNAs, possibly due to haploinsufficiency of this gene.

### 6.1. Data Availability Statement

Resource origin and associated information are described in the key resource table. *Osiris* knock out strains and guide RNA strains created in this study are available from the National Institute of Genetics (https://shigen.nig.ac.jp/fly/nigfly/). Original image stacks of FISH images of each *Osi* gene in .oib format are deposited to the SSBD repository (https://doi.org/10.24631/ssbd.repos.2022.10.256).

## Supporting information

Table S1_Osi expression

Table S2_Osi RNAi

Table S3_KO sequence_gRNAoligo

## Acknowledgements

We thank the Kyoto *Drosophila* Stock Center, National Institute of Genetics, Bloomington *Drosophila* Stock Center, Vienna *Drosophila* Resource Center, Takahiro Chihara (Hiroshima University) and Developmental Studies Hybridoma Bank for providing fly stocks and antibodies. We thank Shin-ichi Ogawa and Yukinori Morita for the support of Helium Ion Microscopy imaging at AIST. We thank Housei Wada for the preparation of the flattened image stack and members of the Hayashi lab for their comments on the manuscript. ZS was supported by the RIKEN Junior Research Associate. This study was supported by a Grant-in-Aid for Scientific Research (19H05548 to SH) from MEXT, RIKEN-AIST Challenge Project Grant to Yukinori Morita, Shin-ichi Ogawa and SH, and JST SPRING (JPMJSP2148 to ZS).

## Legend for Supplemental Materials

### Supplemental Table

**Table S1. Summary of *Osiris* gene expression**

Expression patterns of 16 *Osi* genes in sensory organs and epidermal cells.

**Table S2. Effect of transgenic RNAi experiments**

List of UAS-RNAi strains targeting *Osiris* genes collected from the National Institute of Genetics, Bloomington Stock Center, and Vienna Drosophila Stock Center. Their effect on viability, when crossed to either da-Gal4, actin-Gal4, or neur-Gal4, is shown.

**Table S3. List of Osiris knockout strains and guide RNA sequences**

**Sheet “KO fly series”.**

List of sequenced *Osi* gene alleles. A shaded row of each gene marked WT shows a targeted wild-type sequence. The sequences corresponding to guide RNA are underlined, and the PAM sequences are in bold. For *Osi1, Osi6, and Osi22,* multiple guide RNAs are designed. Only mutants with out-of-frame in/del mutations were saved for further analysis.

**Sheet “gRNA oligo&vector”.**

List of oligonucleotide sequences used to build guide RNA vectors.

## Supplementary Figures

**Figure S1**. Co-staining with anti-phosphotyrosine and anti-Futsch antibodies identifies trichogen and tormogen cells.

Enlarged views of the mechanosensory organs in An3. Anti-Futsch (22C10) antibody (yellow) strongly labeled the shaft part of trichogen and weakly the cytoplasm of soma. Phosphotyrosine (pY) antibody (green) labels the cell outlines. Those markers allowed the identification of *Osi11* expression in trichogen and *Osi12* in tormogen, and were used throughout this study.

**Figure S2.** mRNA FISH patterns of nine *Osi* genes in developing pupal head (42 hours APF).

Representative images of 9 *Osi* gene expression patterns that were judged to be undetectable. Some red signals are nonspecific reactions to the pupal cuticle remnants. The tissue outlines were marked with DAPI staining (cyan).

**Figure S3.** Additional expression patterns of *Osi* genes.

Red: FISH signals of *Osi* RNAs, green: anti-phosphotyrosine, yellow: anti-Futsch (22C10) staining, cyan: DAPI.

***Osiris1.*** Expression was detected in the basal cylinder and further distal cells of the arista. In the eye, it was detected in primary pigment cells and unidentified cells below photoreceptor cells in the ommatidia.

***Osiris3.*** Expression was detected broadly in the epidermis. It was also expressed in tormogen of Mp, Lab and An3.

***Osiris4.*** Two types of expression were observed in the pupal head sampled at 42 hours APF. In the whole head views, the example in left shows the expression in the eye, mechanosensory organ, and gustatory organ (same one shown in Figure 1). Second example shows expression in pseudotrachea in the labella. High magnification views of Mp showed expression in mechanosensory trichogen only in one case, and in both trichogen and tormogen in another case. The example of Lab shows expression in gustatory tormogen. In the example of An2, tormogen expression was strong, but weak trichogen expression was also detected. In the eye, expressions in primary pigment cells and tormogen and trichogen cells of interommatidial mechanosensory bristles were observed.

***Osiris5.*** This gene was specifically expressed in An3 (middle: projection of anterior surface, right: single slice) in a pattern enriched in the bottom-lateral territory. *Osi5* was not detected in Mp.

***Osiris6.*** Expression was detected in cells adjacent to pseudotrachea in Lab (middle) and in primary pigment cells in the compound eye.

***Osiris7.*** Broad expression in epidermal cells was detected. *Osi7* was also expressed in tormogen cells of mechanosensory organs of An2 (top middle: projection view, top right: single slice), Eye (interommatidial bristle cells: lower 5.4 µm deep slice) and Mp (lower right). It is also expressed in primary pigment cells and cone cells of the compound eye (lower left, Figure 4) and pseudotracheal cells of Lab.

***Osiris8.*** Expressed in trichogen cells of mechanosensory organs and gustatory organs. The signal in the eye is the trichogen cells of interommatidial mechanosensory cells. In the arista, *Osi8* was expressed in the dorsal side of the basal cylinder and further distal cells. In Mp, trichogen cells of the mechanosensory organ express *Osi8*.

***Osiris9.*** Expressed in the epidermis (whole head, An2, An3, Mp), primary pigment cells, and unidentified cells (7.56 µm deep section) of the eye. In the Labium, *Osi9* expression in epidermal cells is low or absent.

***Osiris11.*** Expressed in the trichogen cells of mechanosensory and gustatory organs. It is also expressed in the arista.

***Osiris12.*** Expressed in the tormogen cells of mechanosensory (Eye, Mp, An2), gustatory (Lab), and olfactory (An2) organs. It is also expressed in a subset of the epidermis of Lab (medial sections) and An3, and arista cells forming a distal ring of basal cylinder.

***Osiris13.*** Expressed in the trichogen cells of olfactory organs in An3 and Mp. Those are sensilla basiconica (sb) of An3 and Mp. The expression in An3 includes sensilla trichordia (st) and likely sensilla coeloconica (sc).

***Osiris16.*** Expressed in a small subset of the trichogen cells of An3, but not in Mp. The expression is likely to be in sensilla coeloconica (sc).

***Osiris21.*** Expressed in the trichogen cells of all mechanosensory organs (examples in An2 and Mp are shown). It is also expressed in the gustatory organs (Figure 3).

***Osiris22.*** Expressed broadly in the epidermis. Examples of An2 and An3 (two focal plains), Mp are and Lab shown. In arista, *Osi22* is expressed in the dorsal half of the basal cylinder.

***Osiris23* (*gore-tex*).** Expressed in the trichogen cells of sb in An3 and Mp. Weaker expression was detected in the bottom-medial part, likely corresponding to st (Figure 6).

***Osiris24.*** Expressed in the trichogen cells of mechanosensory (An2) and olfactory (An3, Mp) organs. Weak signals in the epidermis are non-specific.

**Figure S4. Co-labeling of *Osi13* (red) and *Osi23* (green) in maxillary palp.** Expression of the two genes mostly overlap, with some differences (cell b in the middle panels).

Figure S5. *Osi17* KD and KO phenotypes in adult and third instar imaginal discs.

A. Wing expansion defect of *Osi17* RNAi (VDRC37457) induced by actin-Gal4. Treated animals also showed kinked leg phenotype (enlarged view with an asterisk).

B. Wing expansion defect in animals with homozygous *Osi17 knock*-*out* mutant clones induced by wing pouch-specific Ubx-FLP recombination.

C. Expression of *Osi17* RNAi in the trachea did not cause wing defect.

D. Wing, leg, and haltere imaginal discs of third instar larvae. GFP marker for wild-type chromosome (green) and F-actin (phalloidin, red). Mosaicisms were induced in the pouch regions of the wing and haltere discs. No mosaicism was detected in the notum, trachea (tr), air sac primordium (as), and adult muscle precursors (amp).

## Supplementary Movies

**Movie S1. Serial cross-sectional views of *Osi23* expression in An3.**

A confocal stack of An3 labeled for *Osi23*/*gox* RNA (magenta) was computationally flattened, separated into lateral (top) and medial (bottom) halves, and presented as a series of sections moving from the surface to the interior.

## References

Ando T., S. Sekine, S. Inagaki, K. Misaki, L. Badel, et al., 2019b Nanopore Formation in the Cuticle of an Insect Olfactory Sensillum. Current Biology 29: 1512–1520.e6. 10.1016/J.CUB.2019.03.043/ATTACHMENT/57EF20BD-A166-47F5-AB37-EDE5C0ACD2F6/MMC5.XLSX

Bellaïche Y., M. Gho, J. A. Kaltschmidt, A. H. Brand, and F. Schweisguth, 2001 Frizzled regulates localization of cell-fate determinants and mitotic spindle rotation during asymmetric cell division. Nat Cell Biol 3: 50–57. 10.1038/35050558

Bhushan B., 2009 Biomimetics: lessons from nature-an overview. Phil. Trans. R. Soc. A 367: 1445–1486. 10.1098/rsta.2009.0011

Blagodatski A., A. Sergeev, M. Kryuchkov, Y. Lopatina, and V. L. Katanaev, 2015 Diverse set of Turing nanopatterns coat corneae across insect lineages. Proc Natl Acad Sci U S A 112: 10750–10755. 10.1073/pnas.1505748112

Brown J. B., N. Boley, R. Eisman, G. E. May, M. H. Stoiber, et al., 2014 Diversity and dynamics of the Drosophila transcriptome. Nature 2014 512:7515 512: 393–399. 10.1038/nature12962

Chai P. C., S. Cruchet, L. Wigger, and R. Benton, 2019 Sensory neuron lineage mapping and manipulation in the Drosophila olfactory system. Nat Commun 10. 10.1038/s41467-019-08345-4

Dietzl G., D. Chen, F. Schnorrer, K.-C. Su, Y. Barinova, et al., 2007 A genome-wide transgenic RNAi library for conditional gene inactivation in Drosophila. Nature 448: 151–156. 10.1038/nature05954

Graveley B. R., A. N. Brooks, J. W. Carlson, M. O. Duff, J. M. Landolin, et al., 2011 The developmental transcriptome of Drosophila melanogaster. Nature 471: 473–479. 10.1038/nature09715

Hartenstein V., and J. W. Posakony, 1989 Development of adult sensilla on the wing and notum of Drosophila melanogaster. Development 107: 389–405. 10.1242/dev.107.2.389

Hunger T., and R. A. Steinbrecht, 1998 Functional morphology of a double-walled multiporous olfactory sensillum: The sensillum coeloconicum of Bombyx mori (Insecta, Lepidoptera). Tissue Cell 30: 14–29. 10.1016/S0040-8166(98)80003-7

Inagaki S., K. Numata, T. Kondo, M. Tomita, K. Yasuda, et al., 2005 Identification and expression analysis of putative mRNA-like non-coding RNA in Drosophila. Genes to Cells 10: 1163–1173. 10.1111/j.1365-2443.2005.00910.x

Kondo S., and R. Ueda, 2013 Highly Improved gene targeting by germline-specific Cas9 expression in Drosophila. Genetics 195: 715–721. 10.1534/genetics.113.156737

Kryuchkov M., V. L. Katanaev, G. A. Enin, A. Sergeev, A. A. Timchenko, et al., 2011 Analysis of Micro- and Nano-Structures of the Corneal Surface of Drosophila and Its Mutants by Atomic Force Microscopy and Optical Diffraction. PLoS One 6: e22237. 10.1371/journal.pone.0022237

Kryuchkov M., O. Bilousov, J. Lehmann, M. Fiebig, and V. L. Katanaev, 2020 Reverse and forward engineering of Drosophila corneal nanocoatings. Nature 585: 383–389. 10.1038/s41586-020-2707-9

Larkin A., S. J. Marygold, G. Antonazzo, H. Attrill, G. dos Santos, et al., 2021 FlyBase: updates to the Drosophila melanogaster knowledge base. Nucleic Acids Res 49: D899–D907. 10.1093/NAR/GKAA1026

Lécuyer E., N. Parthasarathy, and H. M. Krause, 2008 Fluorescent In Situ Hybridization Protocols in Drosophila Embryos and Tissues, pp. 289–302 in Methods Mol Biol.,.

Lees AD, and Picken L. E. R., 1945 Shape in relation to fine structure in the bristles of Drosophila melanogaster. Proc R Soc Lond B Biol Sci 132: 396–423. 10.1098/rspb.1945.0004

Lindsley D. L., L. Sandler, B. S. Baker, A. T. C. Carpenter, R. E. Denell, et al., 1972 SEGMENTAL ANEUPLOIDY AND THE GENETIC GROSS STRUCTURE OF THE DROSOPHILA GENOME. Genetics 71: 157–184. 10.1093/genetics/71.1.157

Ni J.-Q., M. Markstein, R. Binari, B. Pfeiffer, L.-P. Liu, et al., 2008 Vector and parameters for targeted transgenic RNA interference in Drosophila melanogaster. Nat Methods 5: 49–51. 10.1038/nmeth1146

Scalzotto M., R. Ng, S. Cruchet, M. Saina, J. Armida, et al., 2022 Pheromone sensing in Drosophila requires support cell-expressed Osiris 8. BMC Biol 20: 230. 10.1186/s12915-022-01425-w

Scholl A., Y. Yang, P. McBride, K. Irwin, and L. Jiang, 2018 Tracheal expression of Osiris gene family in Drosophila. Gene Expression Patterns 28: 87–94. 10.1016/j.gep.2018.03.001

Scholl A., I. Ndoja, N. Dhakal, D. Morante, A. Ivan, et al., 2023 The Osiris family genes function as novel regulators of the tube maturation process in the Drosophila trachea. PLoS Genet 19: e1010571. 10.1371/JOURNAL.PGEN.1010571

Shah N., D. R. Dorer, E. N. Moriyama, and A. C. Christensen, 2012 Evolution of a large, conserved, and syntenic gene family in insects. G3: Genes, Genomes, Genetics 2: 313–319. 10.1534/g3.111.001412

Shanbhag S. R., B. Müller, and R. A. Steinbrecht, 1999 Atlas of olfactory organs of Drosophila melanogaster 1. Types, external organization, innervation and distribution of olfactory sensilla. Int J Insect Morphol Embryol 28: 377–397. 10.1016/S0020-7322(99)00039-2

Shanbhag S. R., S. K. Park, C. W. Pikielny, and R. A. Steinbrecht, 2001 Gustatory organs of Drosophila melanogaster: fine structure and expression of the putative odorant-binding protein PBPRP2. Cell and Tissue Research 2001 304:3 304: 423–437. 10.1007/S004410100388

Shiga Y., M. Tanaka-Matakatsu, and S. Hayashi, 1996 A nuclear GFP/β-galactosidase fusion protein as a marker for morphogenesis in living Drosophila. Dev Growth Differ 38: 99–106. 10.1046/j.1440-169X.1996.00012.x

Smith C. R., C. Morandin, M. Noureddine, and S. Pant, 2018 Conserved roles of Osiris genes in insect development, polymorphism and protection. J Evol Biol 31: 516–529. 10.1111/jeb.13238

Sobala L. F., and P. N. Adler, 2016 The Gene Expression Program for the Formation of Wing Cuticle in Drosophila. PLoS Genet 12. 10.1371/journal.pgen.1006100

Steinbrecht R. A., 1997 Pore structures in insect olfactory sensilla: A review of data and concepts. Int J Insect Morphol Embryol 26: 229–245. 10.1016/S0020-7322(97)00024-X

Stocker R. F., 1994 The organization of the chemosensory system in Drosophila melanogaster: a rewiew. Cell Tissue Res 275: 3–26. 10.1007/BF00305372

Tilney L. G., M. S. Tilney, and G. M. Guild, 1995 F actin bundles in Drosophila bristles. I. Two filament cross-links are involved in bundling. J Cell Biol 130: 629–638. 10.1083/jcb.130.3.629

Turing A. M., 1990 The chemical basis of morphogenesis. Bull Math Biol 52: 153–197. 10.1007/BF02459572

Vosshall L. B., and R. F. Stocker, 2007 Molecular architecture of smell and taste in Drosophila. Annu Rev Neurosci 30: 505–533. 10.1146/annurev.neuro.30.051606.094306

Wada H., and S. Hayashi, 2020 Net, skin and flatten, ImageJ plugin tool for extracting surface profiles from curved 3D objects. MicroPubl Biol 2020. 10.17912/micropub.biology.000292

Wigglesworth B. Y. V. B., 1948 The insect cuticle

Wodarz A., U. Hinz, M. Engelbert, and E. Knust, 1995 Expression of Crumbs Confers Apical Character on Plasma Membrane Domains of Ectodermal Epithelia of Drosophila.

