## Supplementary figures and images for "*Osiris* gene family defines the cuticle nano-patterns of *Drosophila*"

### Supplemental Figure S1-5

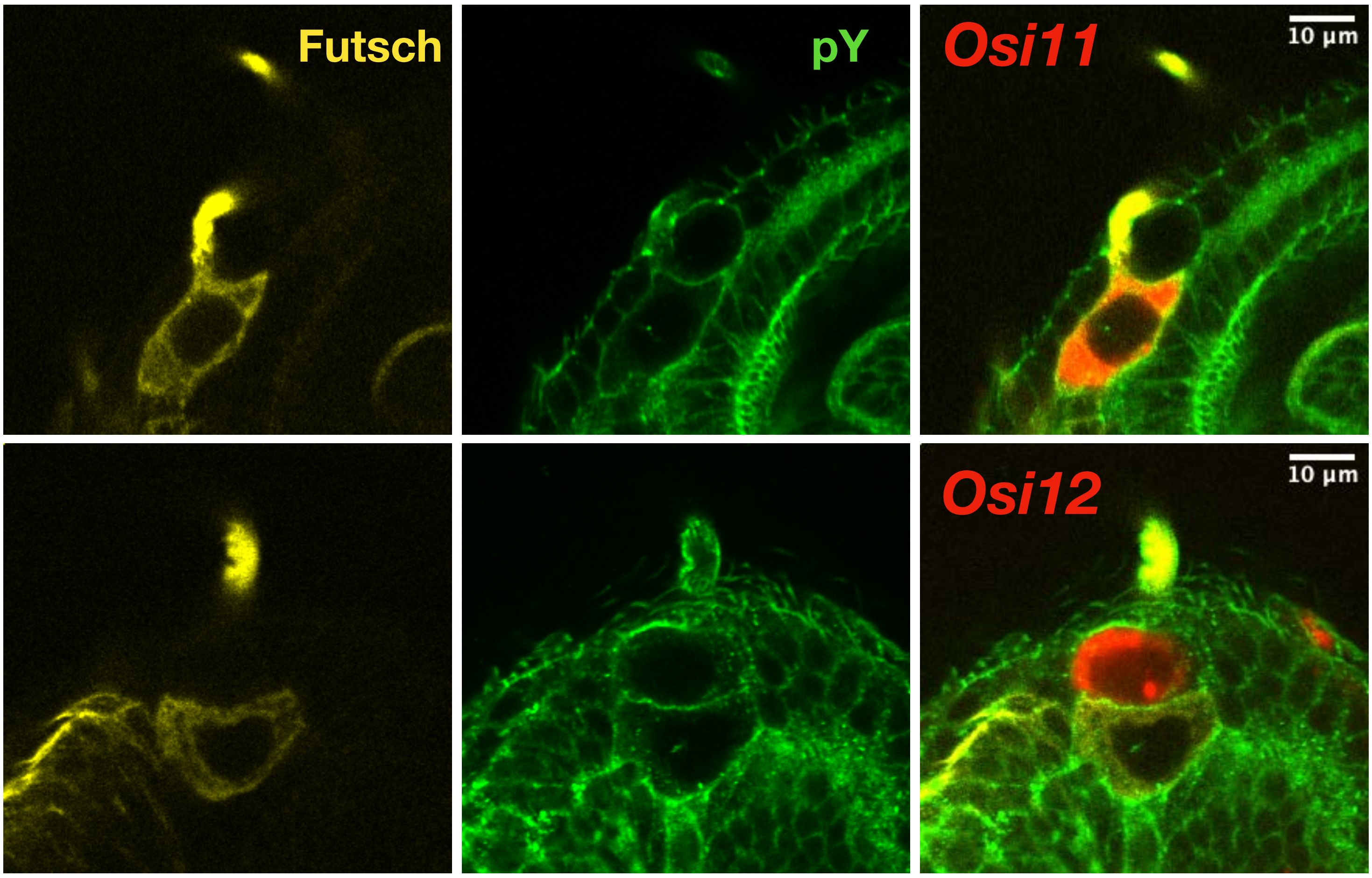


**Figure S1.**

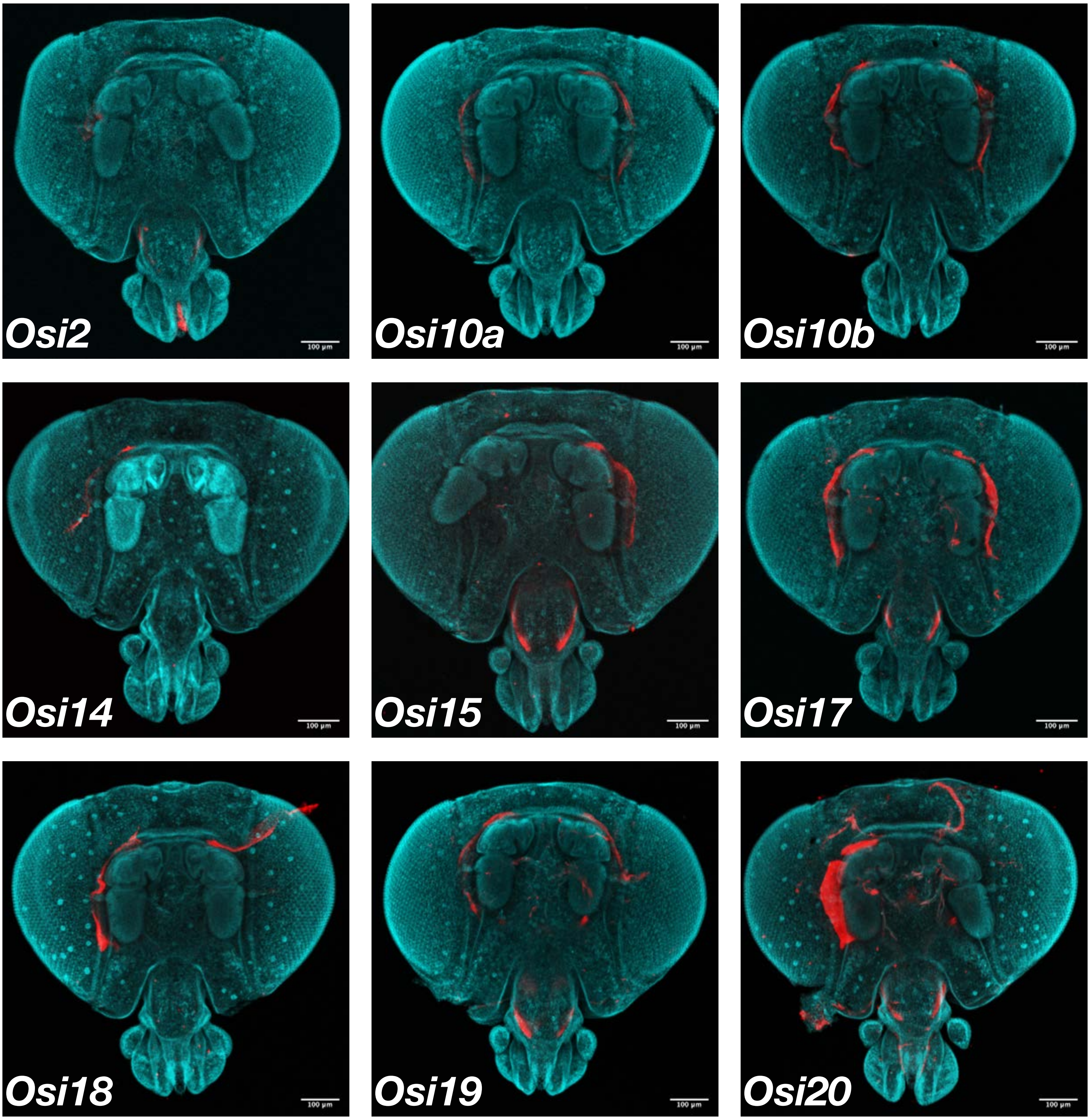


**Figure S2.**

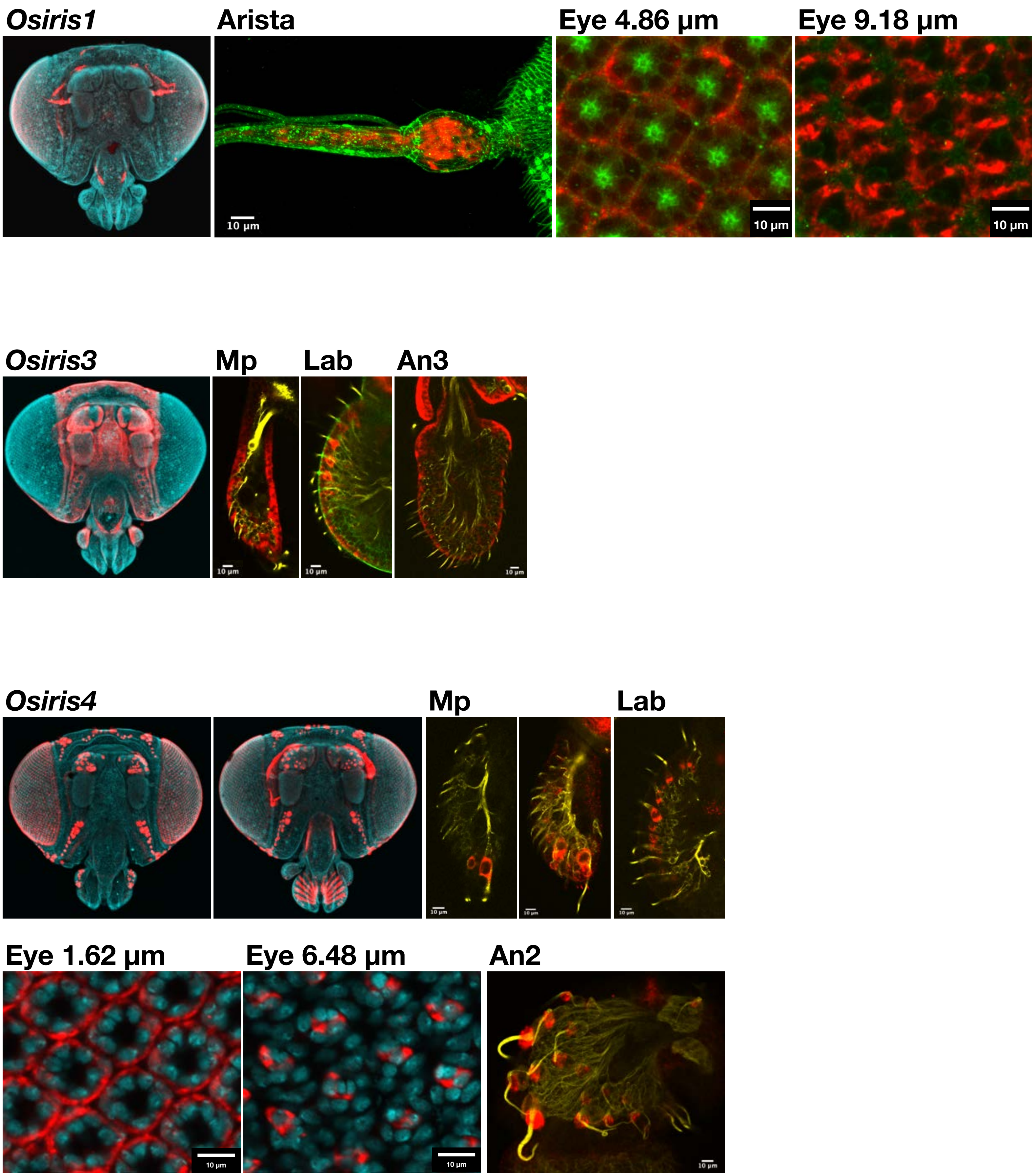

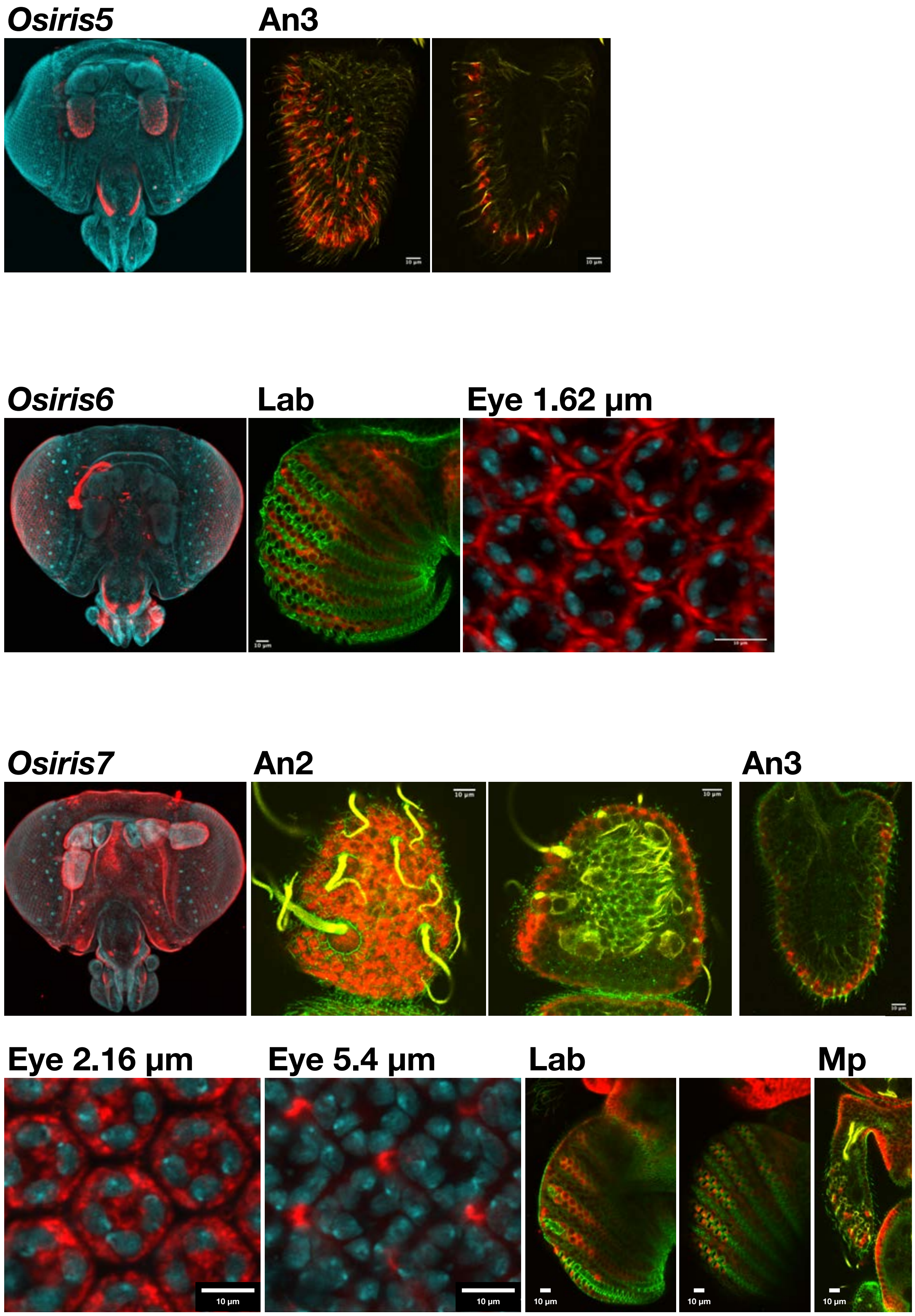

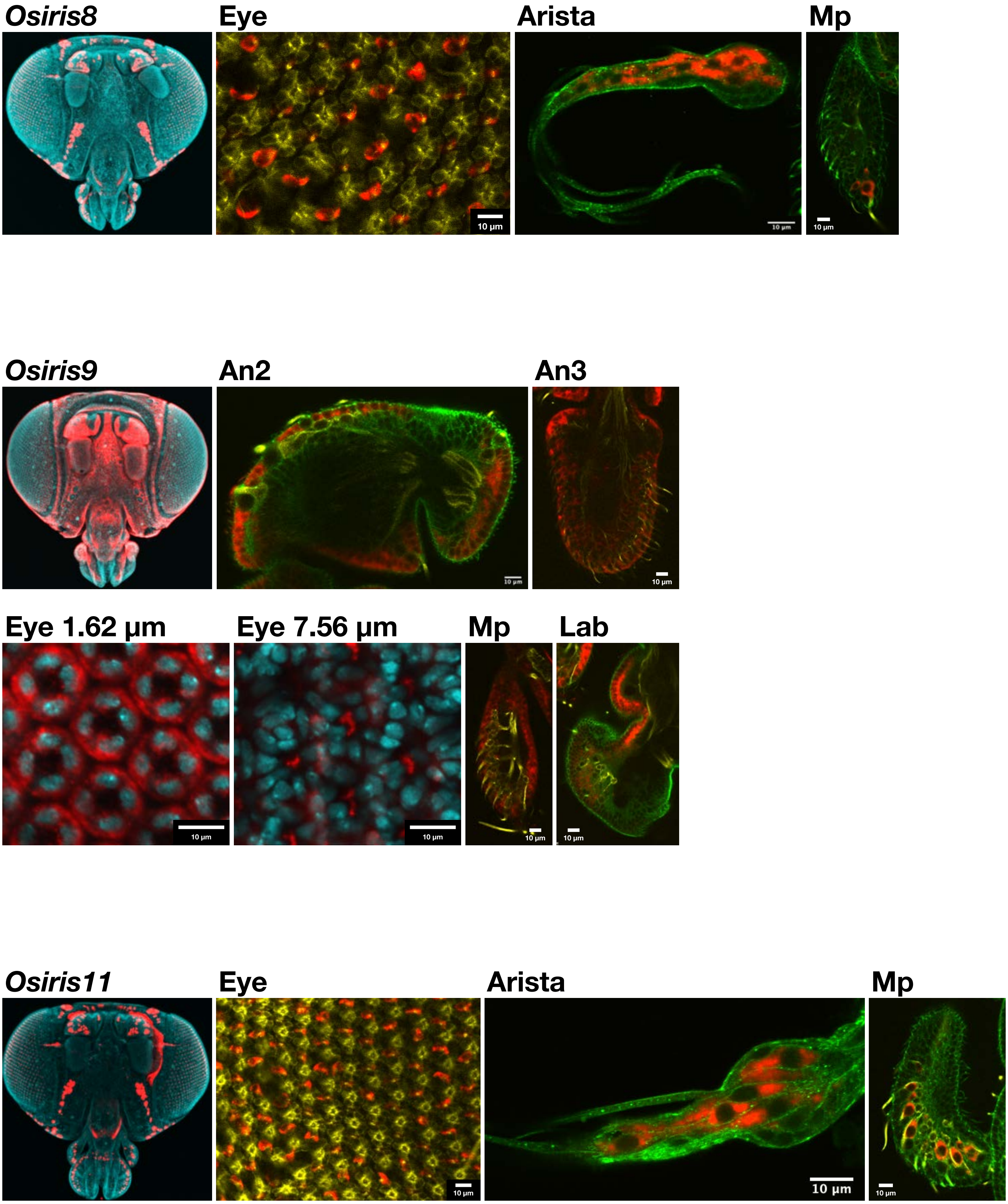

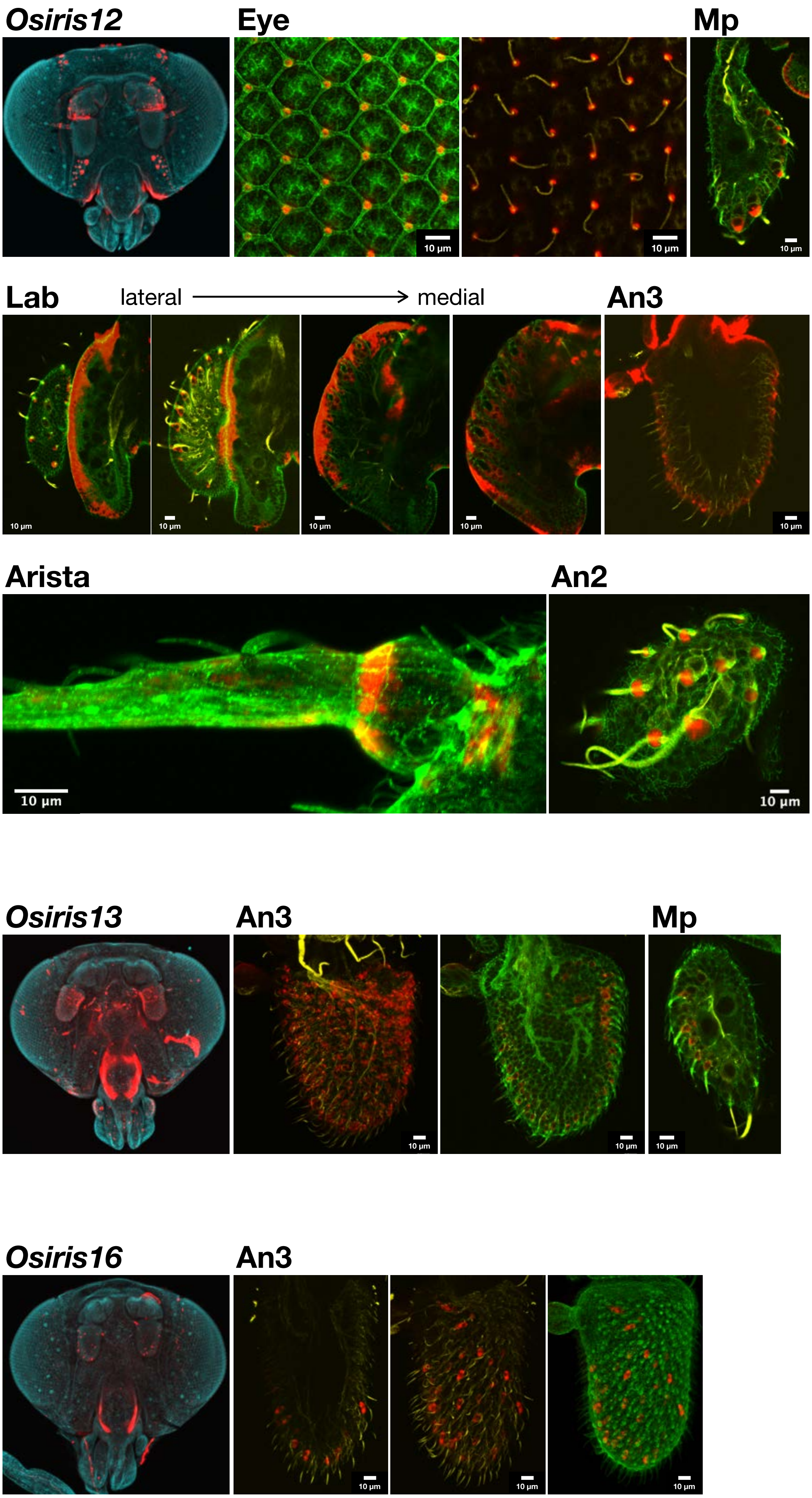

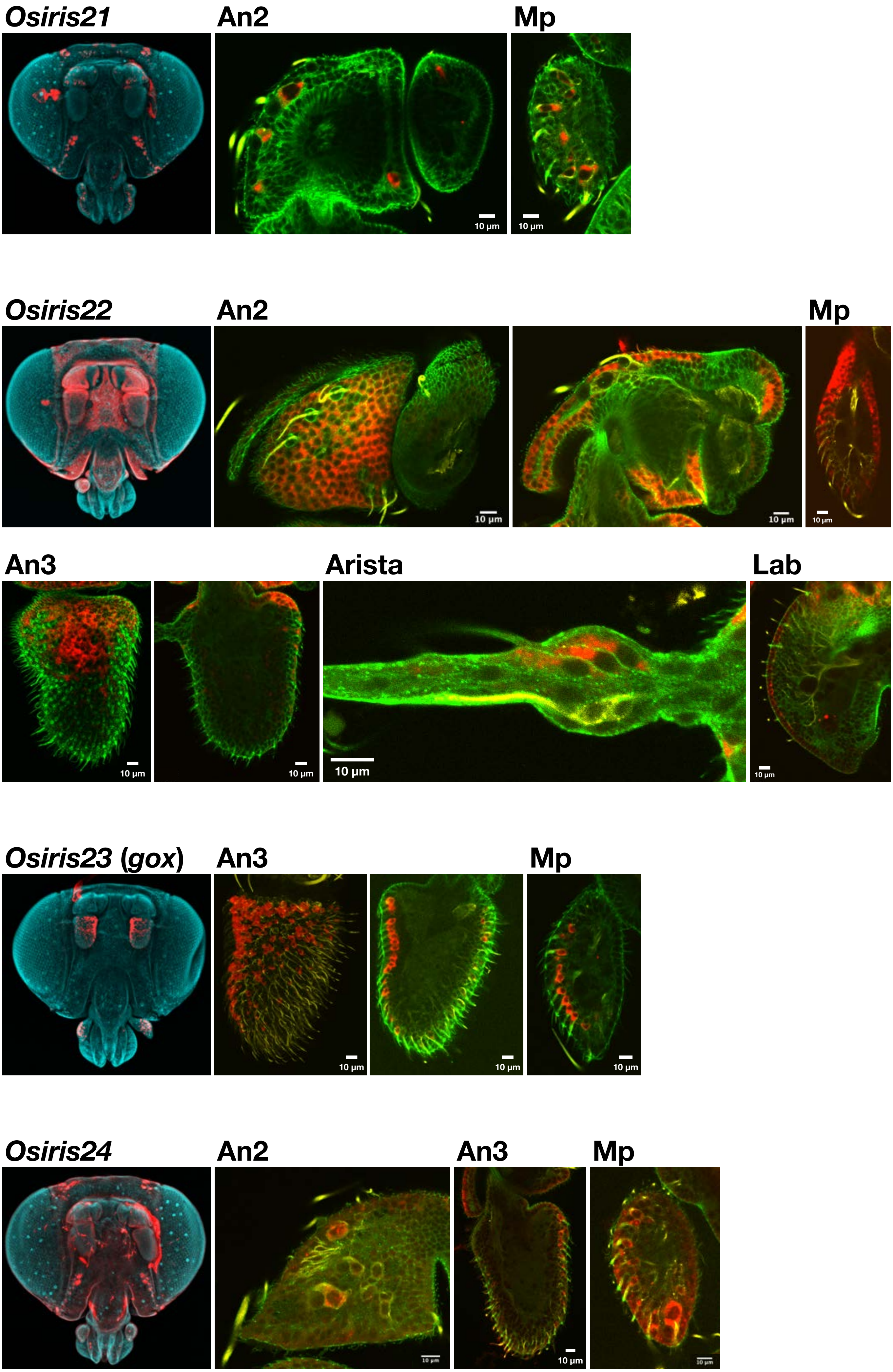

**Figure S4.**

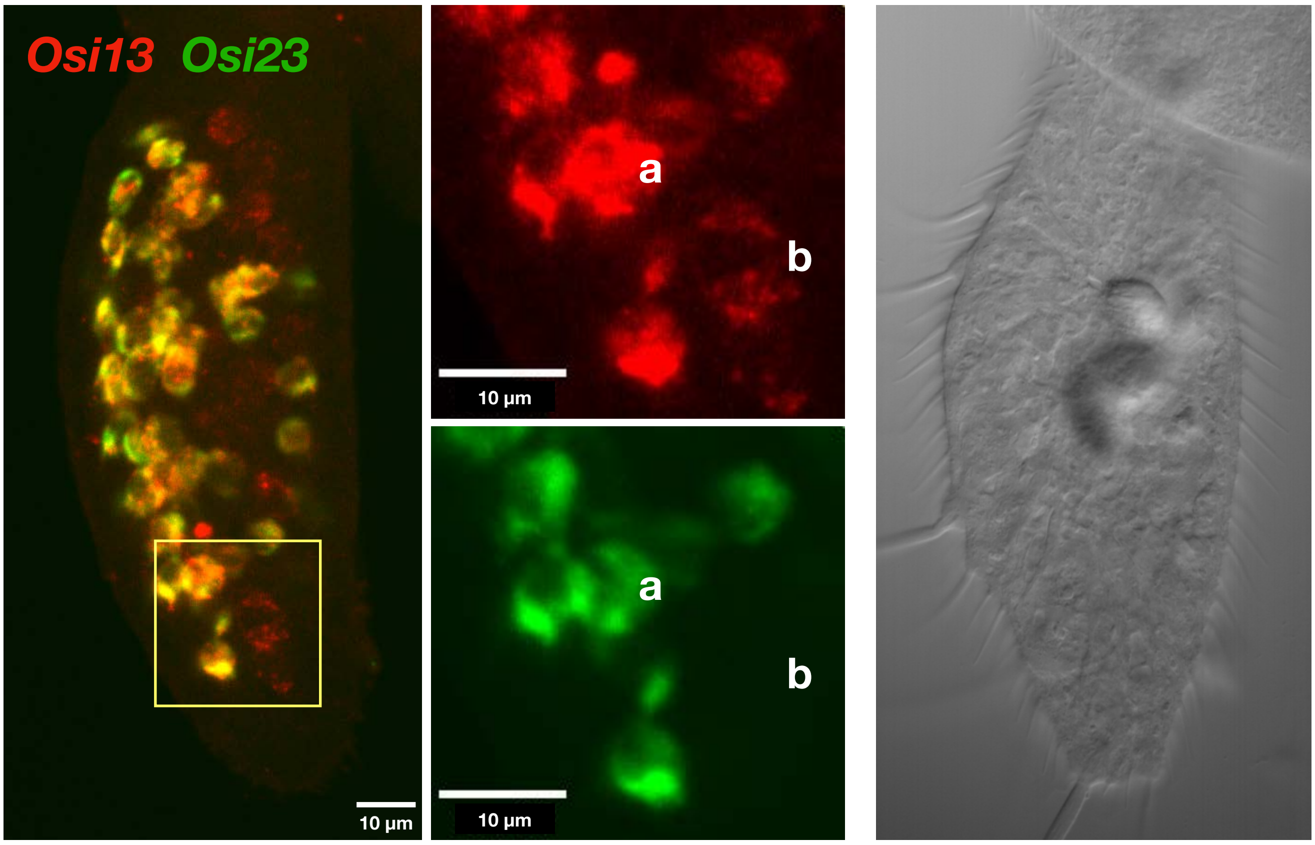

#### Figure S5.

##### *Osiris17*

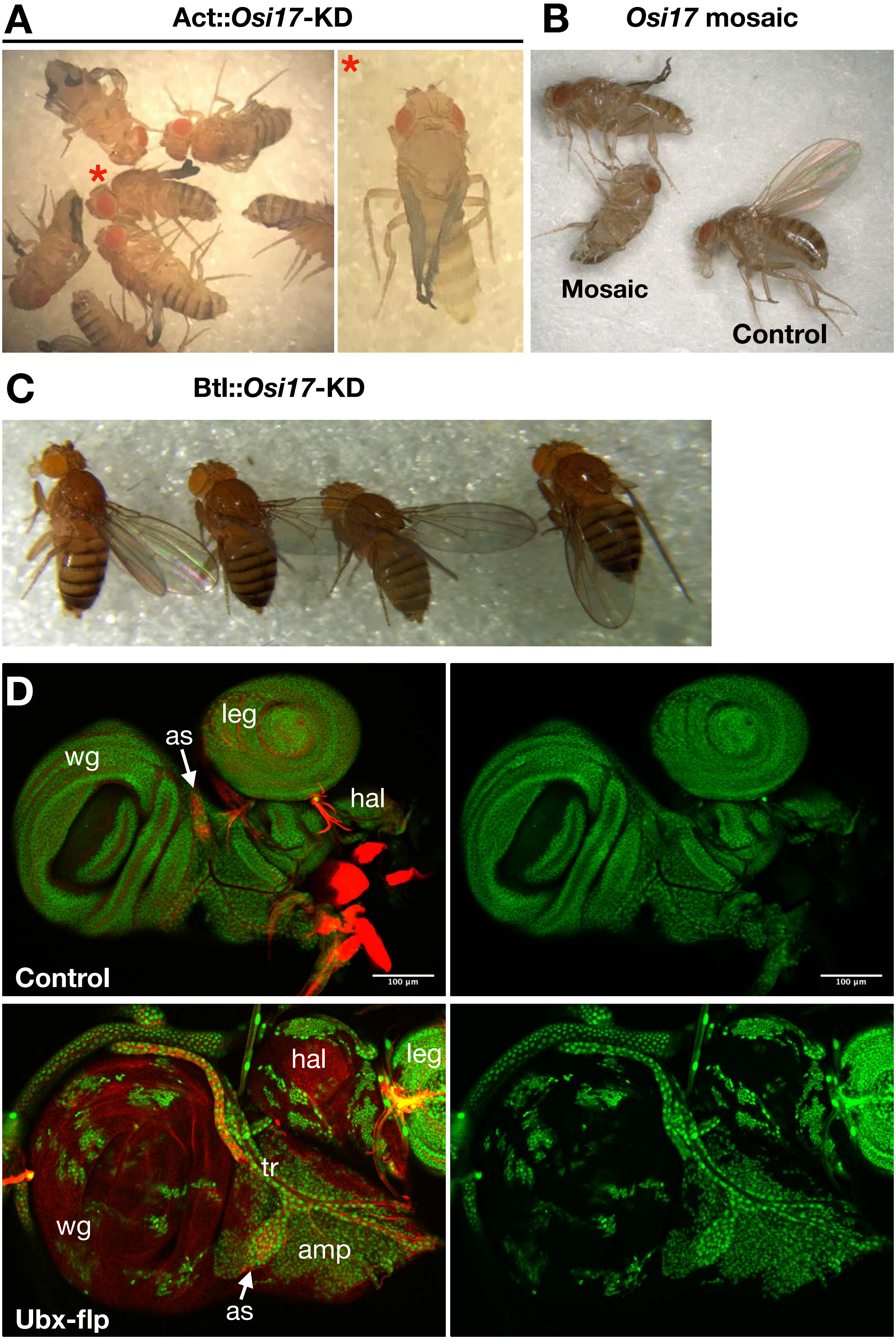
